# APRILE: Exploring the Molecular Mechanisms of Drug Side Effects with Explainable Graph Neural Networks

**DOI:** 10.1101/2021.07.02.450937

**Authors:** Hao Xu, Shengqi Sang, Herbert Yao, Alexandra I. Herghelegiu, Haiping Lu, James T. Yurkovich, Laurence Yang

## Abstract

The majority of people over the age of 65 take two or more medications. While many individual drug side effects are known, polypharmacy side effects due to novel drug combinations poses great risk. Here, we present APRILE: an explainable artificial intelligence (XAI) framework that uses graph neural networks to explore the molecular mechanisms underlying polypharmacy side effects. Given a list of side effects and the pairs of drugs causing them, APRILE identifies a set of proteins (drug targets or non-targets) and associated Gene Ontology (GO) terms as mechanistic ‘explanations’ of associated side effects. Using APRILE, we generate such explanations for 843,318 (learned) and 93,966 (novel) side effect–drug pair events, spanning 861 side effects (472 diseases, 485 symptoms and 9 mental disorders) and 20 disease cate-gories. We show that our two new metrics—pharmacogenomic information utilization and protein-protein interaction information utilization—provide quantitative estimates of mechanism complexity. Explanations were significantly consistent with state of the art disease-gene associations for 232/239 (97%) side effects. Further, APRILE generated new insights into molecular mechanisms of four diverse categories of adverse drug reactions: infection, metabolic diseases, gastrointestinal diseases, and mental disorders, including paradoxical side effects. We demonstrate the viability of discovering polypharmacy side effect mechanisms by training an XAI framework on massive biomedical data. Consequently, it facilitates wider and more reliable use of AI in healthcare.

## Introduction

Polypharmacy poses a growing global health challenge. Polypharmacy is prevalent in the general population, as half of those taking any prescription medication are exposed to two or more drugs^1^. For people 65 and older, this percentage increases to 75%^2^. With 22% of the global population expected to be over 65 years old by 2050^2^, polypharmacy is becoming increasingly pervasive.

Polypharmacy increases the risk of adverse drug reactions (ADRs). In Europe, adverse drug events are responsible for 8.6 million hospital admissions per year^2^. In the UK, up to 11% of hospital admissions were due to adverse drug events—critically, over 70% of these admissions were in elderly patients, who are exposed to polypharmacy^3^. Thus, polypharmacy management is a core element of the global challenge for medication safety that was launched by the World Health Organization (WHO) in 2017^2^.

Polypharmacy management is complex: over 200,000 approved drugs gives rise to over 200 million possible drug pairs, most of which will not be experimentally tested. Thus, Artificial Intelligence (AI)-assisted methods have been developed to predict potential polypharmacy risk^4,5^. Specifically, such “predictors” predict the ADRs for all possible drug combinations.

Pharmacogenomic information plays an essential role for predicting and explaining the drug interactions and ADRs, and there are massive experimental data available^6,7^. This information includes drug targets, functional associations or physical connections between or within drugs and proteins, all of which can be naturally depicted as a knowledge graph (KG). By extracting and encoding such information, graph neural network based deep learning methods are able to provide more reasonable ADR predictions based on the underlying protein interactions^4,5^. However, the prediction accuracy of these approaches is not high enough for the model to be trusted completely and anomalous decisions can be made, causing undesired side effect in practical applications. Thus, human-interpretable explanation for making such predictions are still needed to provide data-driven solutions that help medical practitioners make better diagnoses to improve medical care^8^.

Explainable AI (XAI) seeks to enhance the model training, representation learning and decision-making processes with human-understandable explanations^9^. Recently, increasing numbers of XAI methods have been designed to improve the reliability of model predictions^10,11^. The “explainer” methods have developed as a sub-domain of XAI that explain the predictions made by particular classes of machine/deep learning models, including even those that are non-interpretable^12,13^, such as an explainer for all graph neural networks^14^. We hypothesize that the explanations for ADR predictions based on the pharmacogenomic KG can reveal new mechanistic associations between ADRs and drug targets/non-targets. Such associations make it possible to gain further pharmaceutical insights and provide reliable treatment options.

In this paper, we demonstrate that XAI can effectively reveal the mechanisms underlying polypharmacy side effects. We present APRILE (**A**dverse **P**olypharmacy **R**eaction **I**ntelligent **L**earner and **E**xplainer), an explainable framework in the format of predictor-explainer, that leverages pharmacogenomic information for discovering the molecular mechanisms of interactions between drugs. The workflow of APRILE is as follows: first it trains a machine learning model, APRILE-Pred, that is capable of predicting drug combination’s side effects from pharmacogenomic data. Then, given a side effect prediction or a set of side effect predictions, APRILE trains another machine learning model, APRILE-Exp. This model produces a tentative molecular mechanism level explanation to those predictions—it queries APRILE-Pred to identify the most important pharmacogenomic information required to make the predictions.

## Results

Here, we present APRILE, an XAI platform that provides novel drug-drug-ADR triplet predictions and mechanistic explanations for their underlying mechanisms of interaction. In the following sections, we justify the need for APRILE, detail the framework itself and provide validation for the underlying model. We then show that APRILE generates new insights into molecular mechanisms of four diverse categories of adverse drug reactions: infection, metabolic diseases, gastrointestinal diseases, and mental disorders, including paradoxical side effects of APRILE using existing data for drug interactions and mechanisms.

### XAI is required to explain drug interactions

We first asked whether XAI is needed to explain why drug pairs cause a side effect based on the KG used to train APRILE. Our KG comprised 4.6 million drug-drug-ADR triplets, 18,596 drug-protein interactions and 1,431,224 protein-protein interactions (Methods). If most of the ADRs were caused simply by two drugs targeting the same protein, then AI may not be necessary as the side effect mechanism would be trivial. However, only 2.6% of drug-drug-ADR triplets had at least one shared protein target. Therefore, XAI, such as APRILE, is needed to explain the vast majority of these side effects.

### APRILE: an explainable framework for discovering adverse polypharmacy reactions

The APRILE framework is shown in Figure 1 and described in the Methods section. It is composed of two modules, APRILE-Pred and APRILE-Exp, for predicting and explaining adverse drug reactions (ADRs), respectively.

**Figure 1.**
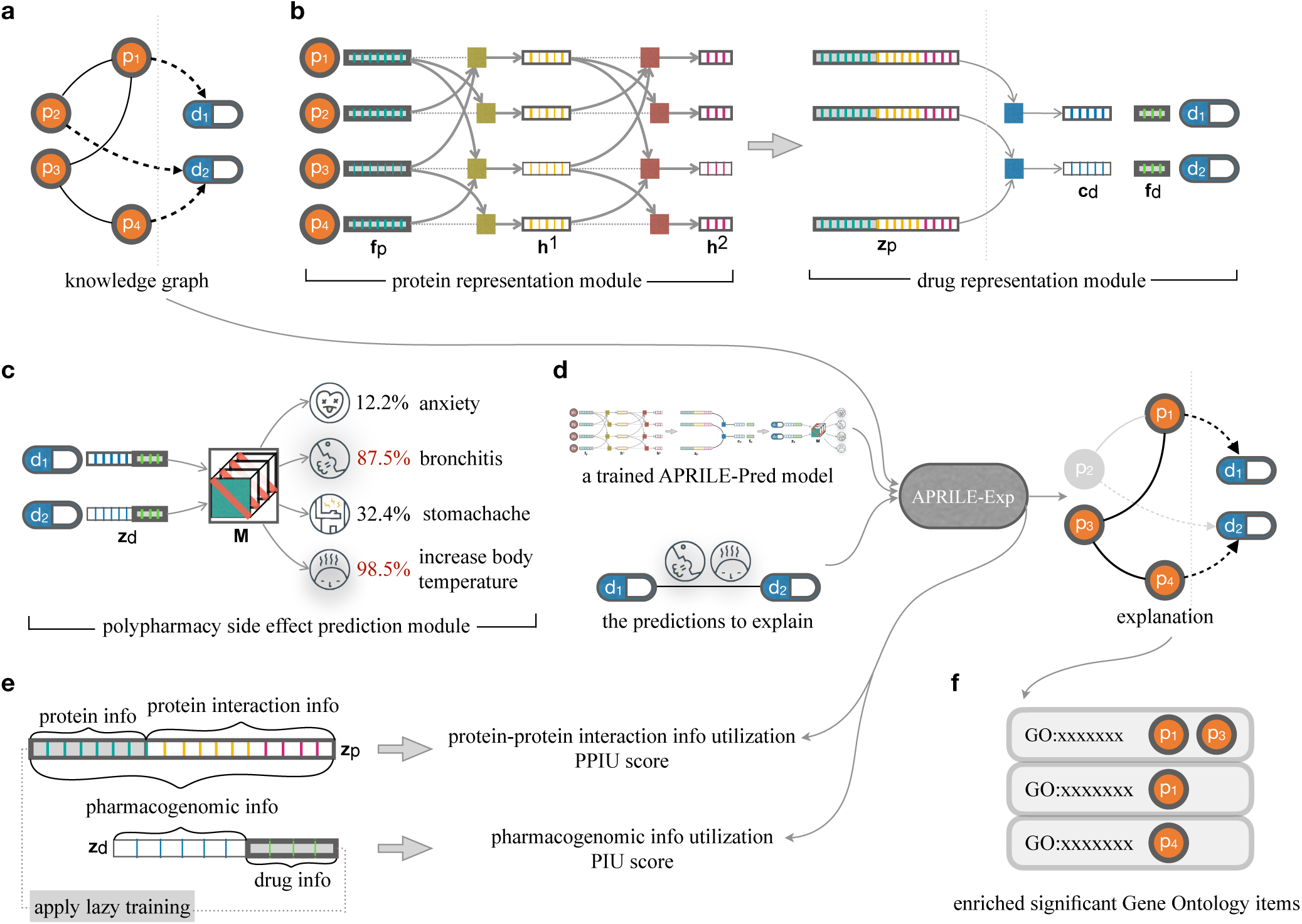
Overview of the APRILE framework: a workflow to predict and explain ADRs. **A-** the pharmacogenomic knowledge graph which contains drug-gene and gene-gene relations. **B-** protein and drug representation modules apply graph convolutional networks to extract pharmacogenomic information from **A**. **C-**, APRILE prediction module uses drug representations to predict the probability of that all drug pairs cause different side effects. The modules in **B** and **C** constitute the predictor of APRILE, APRILE-Pred, and are trained end-to-end with polypharmacy adverse event data. **D-** APRILE-Exp gives a small subgraph of **A** as an explanation for given prediction(s) e.g. polypharmacy adverse event(s), which are the most important drug targets and protein associations in making the given prediction(s), accompanied by functional roles of these proteins. PPIU and PIU score are also provided to measure the utilization of information on drug targets and protein interaction by APRILE-Pred. **E-** lazy training fixes the parameters **f***_p_* and **f***_d_* when they were initialized to prevent the relational information from being learned into the embeddings of protein, drug and side effect. **F-** gene ontology (GO) enrichment analysis is used to investigate the functional roles of the genes in APRILE-Exp’s explanations.

APRILE-Pred is an instance of graph neural network (GNN), which is a popular machine learning model for extracting structural information and node features from graph-like data. GNN has proven to be successful at predicting polypharmacy ADR tasks^4,5^ APRILE-Pred takes a pharmacogenomic KG as input, and predicts the polypharmacy side effects driven by all possible drug pairs. The training of APRILE-Pred is the process of tuning its parameters and learning node embeddings by capturing the information in the pharmacogenomic KG to predict ADRs.

APRILE-Exp is a variant of the GNNExplainer^14^ model, which is a general framework to explain GNNs. To our knowledge, our work is the first implementation of the GNN-explainer to the problem of explaining ADR predictions. However, we find that applying GNN-explainer directly results in explanations that are not biologically informative. Most of the biologically important interactions (protein-protein and protein-drug interactions) are not used for the explanation—instead, only drugs are used to explain the ADR. This is because the drug embeddings learned by the GNN dominate the model explanations. Therefore, we develop a new training strategy for GNNs to enhance explainability, which we call ‘lazy training’ (Methods). This strategy involves tuning parameterization to isolate relational information on the KG. We thus ensure that target and non-target proteins can be considered when explaining ADRs.

Moreover, APRILE-EXP suggests a tailored loss function to provide meaningful and flexible explanations, taking the multiple mechanisms by which polypharmacy can cause ADRs into account. It takes a well-trained APRILE-Pred and a set of drug-drug-ADR triplets as input. Its training amounts to look-ing for a subgraph of the original pharmacogenomic knowledge graph that simultaneously minimizes (i) the number of drug targets and non-targets in the subgraph, and (ii) the error of APRILE-Pred’s prediction(s), which are inputs (Equation (7) in Methods). The competition between minimizing target versus non-target proteins in objective (i) is controlled by the coefficients (α and β) of corresponding terms in the loss function, which can be manually tuned. Finally the optimal subgraph is outputted as a ‘molecular mechanism-level explanation’ of the side effects of interest.

To analyze APRILE’s ADR predictions and explanations, we design two metrics to describe the complexity of ADR mechanisms: pharmacogenomic information utilization (PIU) and protein-protein interaction information utilization (PPIU) scores (Methods). These metrics quantify how important the information on drug targets or non-target proteins is respectively for APRILE-Pred’s predictions, and guide the coefficients tuning in the APRILE-Exp’s loss function. APRILE thus is able to suggest the identification of a given ADR’s mechanism into the following two different classes of mechanisms: mechanisms involving 1) only the drug targets, and 2) target and non-target proteins.

### Lazy training improves the explainability of APRILE

We found that lazy training improves the explainability of APRILE-Pred stably (Figure 2-A,C) without significantly reducing model prediction accuracy. First, we assessed how sensitive the model is to the initialization of these fixed matrices and the size of samples for training in detail (Methods). The results show that the standard deviation of the AUROC score on the corresponding testing set for different splitting rates is 0.038 *±* 0.0004 (Figure 2-A, Table S1). Next, the ablation study on lazy training (Methods) also showed that there is a trade-off between model prediction accuracy and the PPIU score for each side effect (Figure 2-C). Sacrificing a little accuracy can significantly improve the potential interpretation of the model.

**Figure 2.**
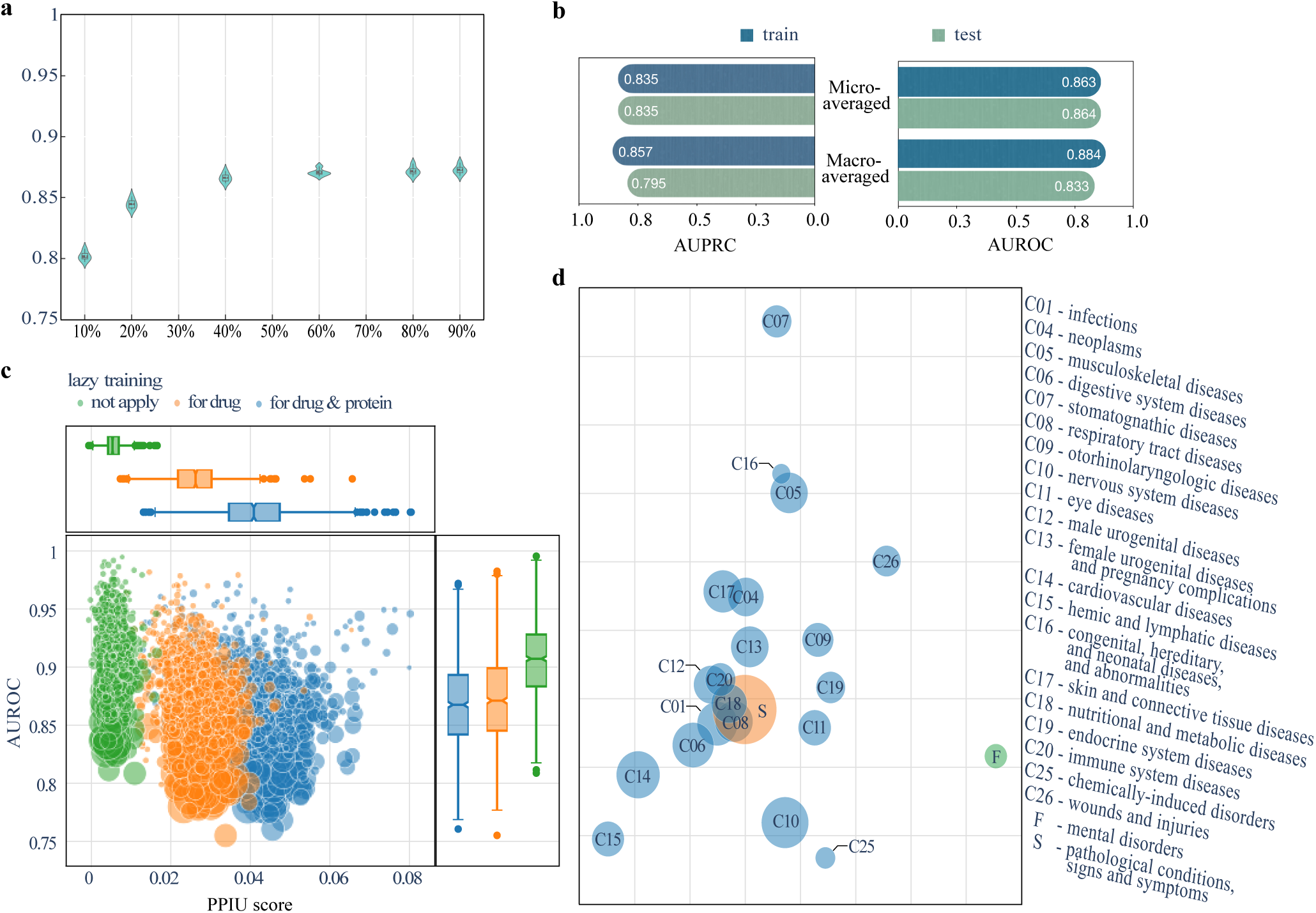
Evaluation metrics and visualization for APRILE-Pred. **A-** Sensitive evaluation for lazy training to different dataset split rates and initial parameter values. **B-** performance comparison of the selected trained APRILE-Pred model on the training and testing sets. **C-** ablation study on lazy training, which compares the distribution of AUROC for each side effect when applying no, partial and full lazy training. Marker size indicates the number of drug pairs that cause the corresponding side effect. **D-** UMAP of the category-average side effect embeddings learned by APRILE-Pred. Marker sizes are the log-scaled numbers of side effects in each categories.

APRILE-Pred model tends to better predict rare side effects than common side effects (the marker size indicates the number of positive instances of the corresponding side effects in Figure 2-C). This observation is as expected because APRILE-Pred emphasizes learning from pharmacogenomic information and weakens learning from drug features and direct drug interactions.

### Validation of APRILE

We built APRILE on a pharmacogenomic knowledge graph involving *∼*1.5 million relations among 284 drugs and *∼*19 thousand drug targets and non-targets. Then, it is trained and tested on *∼*4.6 million drug-drug interactions (DDIs) each associated with an ADR. In order to ensure that APRILE’s performance on the seen and unseen DDIs is balanced, APRILE-Pred’s model selection is based on cross validation. The selected APRILE-Pred model is trained with 90% positive samples, tested with the remaining 10%, and has the most balanced performance on the training and testing set (micro-averaged AUROC=0.85) (Figure 2-B).

### APRILE learns meaningful side effect embeddings

To examine what APRILE-Pred learned, we interrogated the prediction model from side effect to category level and examined its learned embeddings. Based on the Medical Subject Headings (MeSH) database^15^, we classify the side effects into three main categories: *disease* (C), *mental disorder* (F) and *symptom* (S) (Supplementary Data). We found that a two-dimensional UMAP^16^ projection of side effect embeddings learned by the selected trained APRILE-Pred contained meaningful clusters for 20 disease subcategories and the other two categories (Figure 2-D). Clusters F (mental disorder), C07 (stomatognathic disease) and C13 (female urogenital diseases and pregnancy complications) were located at the extremities, indicating that their embeddings differed the most from other MeSH categories and from each other. Clusters close to each other are interpreted as having similar embeddings: e.g., C16 (congenital, hereditary, and neonatal disease and abnormalities) and C05 (musculoskeletal diseases). However, comparing the Euclidean distance and the Jaccard similarity of the side effects of these 22 side effect categories shows that the overlapping side effects among different categories does not dominate the clustering of these categories (Figure S2). Rather, the embeddings are based on extracting information from integrating the pharmacogenomic knowledge graph with drug adverse events.

### APRILE generates valid explanations

When discovering the toxicity mechanisms of combination therapy, APRILE-Exp explains why APRILE-Pred decides that a pair of drugs caused a side effect. We perform a literature-based evaluation of the explanations’ reliability. Literature evidence was found for additional explanations for ADRs in the training set (Table S2). Out of 843,318 predictions (Supplementary Data), we manually searched literature that supported explanations for the top five predictions ranked by prediction score (Methods). We found evidence for 2/3 genes for scotoma, one gene (*CD7*) for melanocytic naevus and none for elevated prostate specific antigen. Indirect evidence was found for bacterial pneumonia and post traumatic stress disorder.

When studying the general mechanism of a side effect, APRILE-Exp takes the set of all predictions on this side effect as input and gives the reasons why APRILE-Pred makes these predictions. For the example of pleural pain, the input to APRILE-Exp is {(drug_1_, drug_2_, pleural pain) *∈* **A***^R^*} (Methods), which includes all drug pairs causing pleural pain. We do this for every side effect (861 in total) and get a set of important genes, gene-gene interactions and enriched GO items (Figure 3-A, Supplementary Data) We found that 6 GO items: ‘GO:0007186’, ‘GO:0007268’, ‘GO:0007204’, ‘GO:0007165’, ‘GO:0006954’, ‘GO:0007200’ are associated with all side effects. Meanwhile, 250 GO items are associated with only one side effect, while there are 159 side effects with unique GO items.

**Figure 3.**
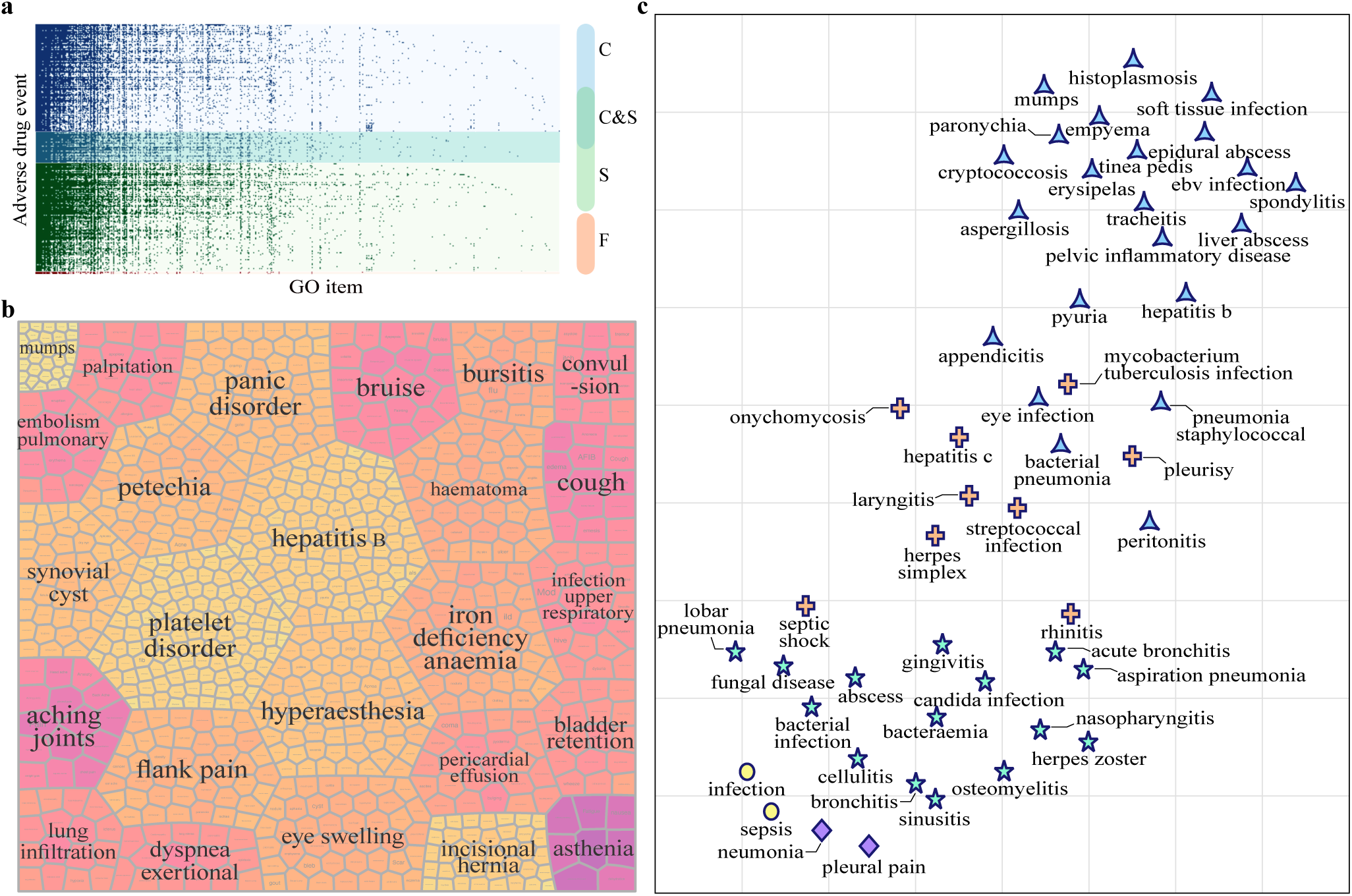
Side effect clustering based on GO items identified by APRILE-Exp. **A-** Association matrix for 861 side effects and 1,787 biological process GO items. C, S, F are side effect categories are the same as those in Fig 2d. **B-** side effect clustering based on the APRILE-Exp’s explanation. The name of the example for each cluster (25 in total) are represented in large fonts. Each small cell is a side effect, and its colour indicts the number of GO items associated with it. (See Table S3 for details). **C-** clustering side effects in the infection disease category based on associated GO items. The side effects in the same cluster use the same marker symbol.

We then investigated the significance of the explanations by comparing them against disease-gene associations from Comparative Toxicogenomics Database (CTD)^17^, for 239 side effects that were found in CTD. We computed the precision and Jaccard index based on comparing the sets of genes in the APRILE-Exp explanation versus the disease-associated genes in CTD.

We then compared APRILE-Exp accuracy with 10,000 random explanations (i.e., gene sets) (Methods). We compared two types of explanations: all genes in a APRILE-Exp subgraph, or the subset of these genes that are in enriched GO terms. Additionally, two types of random models were tested: (A) one that preserves the number of drug target vs. non-target genes, and (B) one that samples from all genes in the KG. Thus, these random explanations capture protein interactions better than the naive GNN-explainer implementation.

Overall, APRILE-Exp explanations for 97% (232/239) of side effects were better than 10,000 random explanations (*P <* 0.05) (Supplementary Data). On average, the precision of APRILE-Exp explanations was 33% to 55% higher than random models, when comparing GO-enriched explanations versus drug target-agnostic random models (random model B).

For the 5 to 7 side effects where *P >* 0.05, the number of disease-associated genes in CTD was typically small with most *<* 60, one case having 629 genes, and one case having 10,620 genes. For cases with very few CTD genes, we propose that APRILE-Exp explanations may contain new hypotheses for side effect-gene associations. In the last case (proctitis), the random models had a mean precision of 0.5. Thus, by randomly sampling genes from the KG, half are expected to be in the disease-associated gene list. This case suggests that proctitis may be highly complex, involving a multitude of genes, which makes a mechanistic explanation particularly challenging.

### ADR classification using identified biological processes

Very few systematic methods are available for performing a global comparison of molecular mechanisms for a vast number of side effects. To address this challenge using APRILE, we constructed an ADR feature matrix based on the 1,787 biological process GO terms in APRILE’s explanations for 861 side effects (Figure 3-A). We applied affinity propagation^18^ to cluster disease and mental disorder ADRs, based on the negative squared Euclidean distances. We then tuned the input preferences to minimize the number of clusters until all clusters contain more than one side effect. Finally, we obtained 25 side effect clusters in total (Figure 3-B, Table S3).

We hypothesized that side effects clustered by biological process may also be related by MeSH disease categories. Enrichment analyses of the 25 clusters for 22 MeSH categories (see Methods) was used to rank order clusters by odds ratio, subject to an unadjusted enrichment *P <* 0.05 (Figure S1). The top 5 enriched clusters and their MeSH categories were panic disorder-mental disorders, embolism pulmonaryhemic and lymphatic diseases, cough-respiratory tract diseases, palpitation-cardiovascular disease, and incisional hernia-eye diseases. While all five clusters had unadjusted *P <* 0.05 and odds ratio*>* 4, their FDR-adjusted *P* values all exceeded 0.05 (Supplementary Data). This result showed that side effects sharing similar mechanisms (i.e., biological processes) are not necessarily similar according to disease categorizations, in this case MeSH. We thus performed GO enrichment (see Methods) of each cluster and found 1 to 124 enriched GO biological processes (FDR-adjusted *P <* 0.05; median odds ratio *>* 4.34 (Supplementary Data). We further identified up to 64 GOs that were uniquely enriched for each cluster. Thus, side effects that are conventionally considered to be distinct disease categories may share many underlying biological processes. This view of side effect clusters may help in developing interventions against multiple, seemingly disparate side effects.

The bursitis cluster includes 20 side effects including breast cancer, sleep apnea, renal cyst, and gastroduodenal ulcers. These side effects span 10 diverse disease categories. Only one GO term was uniquely enriched: aminergic neurotransmitter loading into synaptic vesicle. In the context of breast cancer, drugs that dysregulate aminergic neurotransmitters affect immune function and cancer progression^19^. Clustering of the side effects sleep apnea and muscle disorder was also supported by literature: aminergic neurotransmitter factors or imbalances can cause sleep apnea^20^, or masticatory muscle disorder^21^. Thus, non-intuitive, mechanistic similarities were captured by APRILE-Exp.

The panic disorder cluster includes 54 side effects spanning 19 disease categories, including mental disorders, nervous system diseases, and cardiovascular diseases (Supplementary Data). It is enriched for 11 unique GO terms, including GOs related to regulation of signal transduction, regulation of ion transmembrane transporter activity, tryptophan catabolism, and fatty acid metabolism and nucleoside salvage (Supplementary Data). Consistent with this clustering, tryptophan metabolism is disturbed in cardiovascular disease patients^22^ and also in panic disorder^23^. In both cases, cytokine-based signal transduction alterations cause disrupted tryptophan metabolism.

### Using PPIU scores to interpret APRILE results

We defined the protein-protein interaction information utilization (PPIU) metric to help interpret APRILE-Exp results (Methods). The PPIU score is a measure of the contribution of known protein-protein interactions to an APRILE prediction. A low score indicates that the target proteins are sufficient for explaining a side effect, while a high score indicates that additional proteins in the network, including those connected to the direct drug target, are needed to fully explain the side effect. Thus, we explored the interpretation of PPIU scores and how they can be used to better understand APRILE results.

Selecting sepsis as an example side effect, we examined the high-probability (*>* 0.95) APRILE predictions for which alternative explanations had low or high PPIU scores. Of the 66 predicted explanations with low PPIU (*<* 0.1), 64 (97%) were direct drug targets (Supplementary Data). Alterations in one such direct target, transferrin, has been reported in sepsis patients^24^, suggesting that a perturbation to transferrin through some drug mechanism may increase the chances of experiencing sepsis as a side effect. Predicted explanations with high PPIU (*>* 0.9) included six target genes and one common non-target gene. All seven genes were related to choline transport and anti-flammatory mechanisms, a neuroimmune pathway that may play a role in the development of sepsis.

### APRILE predicts drug combinations with altered individual drug effects

We next explored predicted drug-drug-ADR triplets that were not in the training set, meaning they may represent yet undocumented ADRs. One such novel, high-confidence (prediction score *>* 0.99) example is that the combination of omeprazole and gabapentin were predicted to result in pleural pain. Omeprazole is an H+/K+ ATPase-binding proton-pump inhibitor typically used to treat gastric acid-related disorders, and gabapentin is a voltage-gated calcium channel (VGCC) blocker typically prescribed as an anticonvulsant. For this pair, APRILE predicted significant enrichment for GO terms associated with VGCCs (cellular component ‘VGCC complex’ and molecular function ‘VGCC activity’).

The inhibition of H+/K+ ATPase activity—such as by drugs like omeprazole—disrupts the sodium-calcium balance, leading to an accumulation of intracellular calcium^25^. While VGCC blockers such as gabapentin are typically used to reduce pain by inhibiting the transmission of pain through calcium gradients, VGCC blockers have also been used to reduce intracecullar calcium accumulation^26^. Furthermore, calcium channel blockers have been thought to cause pleural effusion, which can cause pleural pain^27^. Thus, it is possible that the combination of these two drugs results in not only differing mechanisms of the individual drugs, but also pleural pain. While such a prediction warrants further investigation, it nevertheless suggests that APRILE has the potential to identify common drugs that result in unintended side effects when used in combination.

### Predicting explanations for paradoxical side effects

Some drugs have side effects reported in some conditions but not others, or, in more severe cases, one side effect in some cases and the opposite side effect in other cases. Cases where the drug effect is opposite to the expected outcome is referred to as ‘paradoxical adverse reactions’^28^. Such paradoxical side effects offer obvious challenges for patient care. Here, we explore the capacity of APRILE to predict mechanistic explanations for observed paradoxical side effects through two case studies.

Nicotine activates *CHRNA7*, the human alpha 7 nicotinic acetylcholine receptor. It has been shown to both cause and reduce anxiety, while also having been observed to both desensitize and activate nicotinic acetylcholine receptors (nAChRs)^29^. Ondansetron is an antiemetic which is predicted to bind to *CHRFAM7A*^30^, a chimeric gene partially duplicated from *CHRNA7*. APRILE correctly predicted that the combination of nicotine and ondansetron would result in anxiety. The predicted explanation included two genes that are not direct targets of either drug: *Lynx-1* and *SLURP-1*, which are both endogenous allosteric modulators of nAChRs. *SLURP-1* inhibits alpha 7 nAChR activity (ion influx through the receptor channel)^31^, while *Lynx1* inhibits nAChRs through direct binding and reducing their sensitivity to acetylcholine.

Because both drugs activate *CHRNA7*, we hypothesize that polypharmacy of nicotein and ondansetron induces nAChR hypersensitivity, resulting in increased anxiety. This hypothesis has been corroborated by animal studies, wherein hypersensitive nAChR has been shown to increase anxiety in mice^32^. A simple description of the mechanism is that multiple drug-protein and protein-protein activities combine in a new context, resulting in the polypharmacy side effect. Here, we propose that hypersensitivity to polypharmacy arises as the respective drug target proteins also inhibit a common protein that is not directly targeted by either drug (Figure 4b). We explored other predicted cases of paradoxical side effects, such as how the use of a combination of anti-epileptic drugs may present side effects similar to that of an excessive dosage of a single drug (Figure S3)

**Figure 4.**
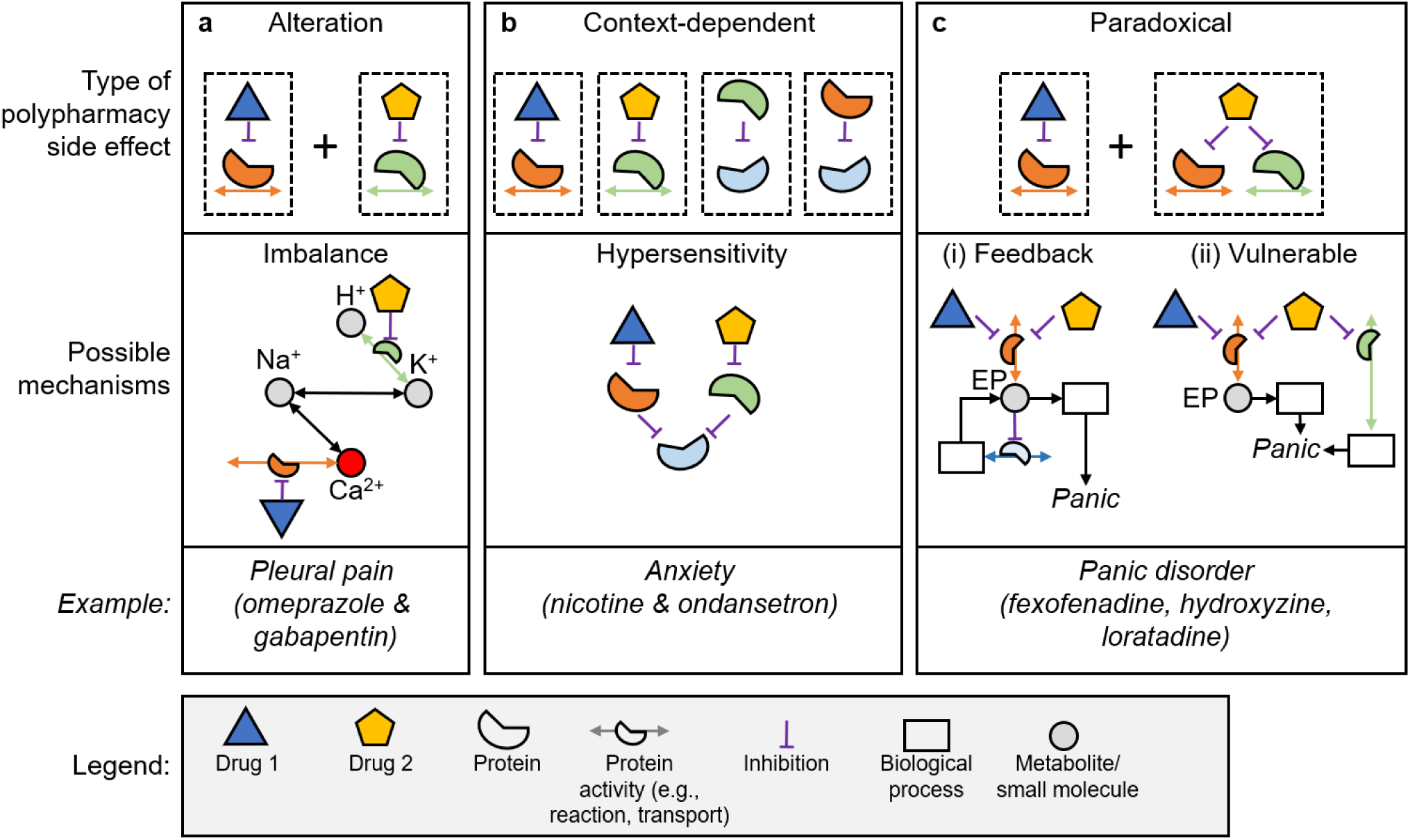
Mechanisms of ADRs in polypharmacy. **A-** polypharmacy alters drug effects resulting in molecular imbalance. **B-** drug effects change with context, such as drug targets inhibiting a common non-target protein. **C-** paradoxical ADRs may be caused by (i) inverse response within feedback regulatory processes, or by increased (ii) vulnerability to disruption of a biological process due to the action of a second drug.

### Suggesting mechanisms for common polypharmacy side effects

Finally, we performed a global analysis to investigate polypharmacy side effects that commonly result from pairs of drugs. Specifically, we examined three common peptic ulcers—duodenal, gastric, and esophageal—that are documented side effects for 69 drug pairs. APRILE was used to predict mechanistic explanations these three side effects have in common, resulting in seven GO terms (Table S4). *Helicobacter pylori* infection is a common cause of ulcers^33^ and is thought to cause ulcers in nearly 20% of human hosts^33^. This process led us to hypothesize two key mechanisms related to *H. pylori* infection in the gut microbiome that duodenal, gastric, and esophageal ulcers share.

First, APRILE predicted heme binding (GO:0020037) and lowered iron binding (GO:0005506) to be common mechanisms in these peptic ulcers. *H. pylori* virulence has previously been linked to both heme-binding proteins^34^ and reduced iron-binding capacity^35^. Second, APRILE predicted that several metabolic processes—arachidonic acid metabolism, geranyl diphosphate biosynthesis, and phospholipase A2 activity (GO:0004623)—contribute to the cause of these ulcers. These processes are known to play a role in breaking or maintaining the mucosal barrier, which helps defend against infection^36^. Thus, while the virulence of the pathogen may contribute, APRILE identified important host factors that increase susceptibility to infection—ultimately increasing the risk for peptic ulcers.

## Discussion

The ability to predict and mechanistically understand drug side effects resulting from polypharmacy poses a significant challenge to modern medicine. Here, we employ a novel explainable artificial intelligence (XAI) platform, APRILE, to provide meaningful, accurate, mechanistic explanations for polypharmacy side effects. APRILE computes quantitative metrics that indicate whether proteins beside drug targets are likely to be involved in an ADR mechanism. We validated APRILE predictions, and offered several mechanistic hypotheses for novel drug-drug-ADR triplets. Taken together, these results have several important implications.

First, our data-driven XAI approach has the potential to identify unreported drug pairs that pose risk to patients and further predict mechanistic explanations to support clinical decision making. There are over 20,000 approved prescription drugs giving rise to over 200 million possible drug pairs. Clinical decision support softwares such as that by uMETHOD Health^37^ create personalized treatment plans that provide physicians with alerts and recommendations on drug interactions. A desired feature of such software is proposing mechanisms that underlie the drug interactions^38^, while also accounting for a patient-specific conditions such as other chronic illnesses (see Supplementary Note 1). Thus, APRILE may be integrated into clinical decision support software to enhance their effectiveness and adoption by clinicians.

Second, lazy training encourages APRILE’s explainability, and APRILE-Exp’s loss design provides meaningful and flexible explanations. Lazy training enables GCNs to maintain the independence of relational information, such that it encourages APRILE-Exp to pay more attention to the non-target proteins that are important for APRILE-Pred to make predictions. We also present the strategy for designing alternative loss functions for the explainer. Users of APRILE can tailor explanations involving drug targets and non-targets by exploring the loss function design space. Therefore, these ideas can help adapt the GNN and GNN-based explainer^14^ to different application scenarios.

Third, an XAI platform that predicts polypharmacy ADRs can help in designing clinical trials by acting as an *in silico* screening step. Robust clinical trials requires selecting a study population while balancing the trial’s capacity to validate the intervention and its generalization to the general population that may deviate from ideal conditions^39^. In particular, comorbidities are common for patients in real world settings, and “pragmatic” clinical trials are increasingly used at or after phase 3 trials^40–42^. Thus, while designing pragmatic clinical trials, APRILE can predict drug-drug interactions that can arise for participants on existing medications and also provide explanatory insights as to why they occur. Overall, this XAI platform for model-driven discovery of novel drug interaction mechanisms could accelerate existing drug-discovery pipelines. Especially for drugs intended to be used in combination therapies (e.g., cancer cocktails) or targeting populations having high polypharmacy (e.g., aging populations).

Using APRILE, we have reported both predicted polypharmacy drug-drug-ADR triplets and predicted mechanistic explanations underlying them. While some of the predictions reported here were corroborated by literature evidence, many require follow-up experiments by the biomedical community before being able to evaluate their clinical utility. In fact, APRILE has the capacity to generate model-driven hypotheses for mechanisms underlying 34 million drug-drug-ADR combinations using existing data—this hypothesis space will grow exponentially as the knowledge graph is expanded with new data. Validating these hypotheses may uncover new drug interaction mechanisms that have translational importance and clinical value.

Here, we present a first step toward AI-guided polypharmacy side effect explanation. Using massive data that already exists, APRILE has generated numerous model-driven hypotheses for ADR mechanisms. APRILE’s high explainability makes it useful when designing safe polypharmacy interventions, whose side effects are too complex to rely on human analysis alone. XAI such as APRILE will play an increasingly critical role for polypharmacy management in an aging population for which polypharmacy is currently the norm. Further, we believe APRILE represents an important foundation for future AI-based platforms that will aid medical practitioners in clinical diagnoses and decision making.

## Methods

### Pharmacogenomic knowledge graph construction

We use two subdatasets for polypharmacy-associated ADR modelling from the BioSNAP-Decagon dataset^7^ to prepare the pharmacogenomic knowledge graph : **PP (protein-protein)**: relations between proteins depend on whether they have functional association or physical connections. **GhG (drug-protein)**: relations between drugs and their targeted proteins. All drugs were referenced by CID, which is linked to the chemical structure; this annotation ensures uniqueness within the database.

In summary, the pharmacogenomic knowledge graph contains 19,365 vertices (284 drugs (*D*) and 19,081 proteins (*P*)), 1,449,820 edges. Among the edges, there are 1,431,224 protein interaction edges (P-P) and 18,596 drug protein targeting edges (P-D). We use the following data structures to represent the graph we constructed.

- **A***^p^ ∈* R*^|P|×|P|^*: symmetric adjacency matrix for the undirected subgraph within proteins.
- **A***^t^ ∈* R*^|P|×|D|^*: adjacency matrix for the directed bipartite subgraph between the proteins and drugs, whose edges are directed from proteins to drugs.

### APRILE-Pred: explainable adverse polypharmacy reaction prediction with GNNs

APRILE-Pred is a Graph Neural Network (GNN) model that takes protein-protein and protein-drug interaction graphs as input and predicts the side-effects caused by drug combinations. Specifically, we first learn the protein representation in the protein interaction subgraph, and then generate the drug representation through the protein-drug interaction subgraph with drug attributes (See Algorithm 1). In addition, The input drug/protein node features/attributes *{***v***_d/p_|d ∈ D, p ∈ P}* are set up to one-hot vectors, which are binary vectors representing the *i*th drug/protein if its *i*th element is 1. Protein and drugs are indexed respectively.

### Learning Drug Representation

Firstly, we use a 2-layer graph convolutional network (GCN)^43^ to capture protein attributes and relations on the protein interaction subgraph **A***^p^*. The input is the protein node features 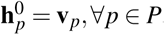. For each protein node *p ∈ P*, the relation between two hidden layers is given by

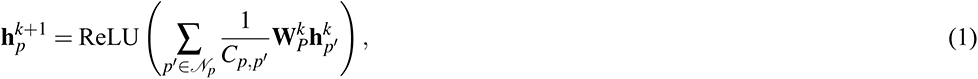

where 𝒩*_p_* denotes the union of protein node *p* and its neighbors on the subgraph 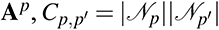 is coefficient calculated by the product of the degree of *p* and *p^′^ ∈ 𝒩_p_*, and **W***_P_*s are parameters. The final learned protein embeddings/representation 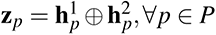 are the concatenation of the outputs of all the GCN layers.

Then, we use a graph-to-graph information propagation module^5^ to transform learned protein embeddings and drug node features into feature representation of drugs via the bipartite protein-drug integration subgraph **A***^t^*. The protein’s contribution to the drug *d*’s representation for all *d ∈ D* is:

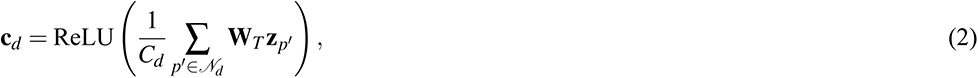

where 𝒩*_d_* = {*p^′^| p^′^ ∈ P,* **A**_p′,d_ = 1} is the set of drug *d*’s protein neighbours, *C_d_* = *|𝒩_d_|*, and **W***_T_* is a trainable parameter matrix. Besides, A linear transformation followed by an activation function is used to map the original drug node features **v***_d_* into the same space that protein’s contributions **c***_d_* live in for each drug *d ∈ D*:

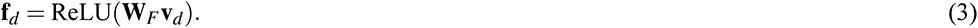

The final representation of drug *d ∈ D* is: **z***_d_* = **z***_p_ ⊕* **c***_d_ ⊕* **f***_d_*, where *⊕* denotes concatenation.

### Polypharmacy Side Effect Prediction

We predict if drug combinations can cause any side effect. Here, DistMult factorization^44^ is used as the scoring function, because it is a well-known model that models rich relationships well on standard benchmarks alone or as a decoder. The probability that drug pairs (*d, d^′^*)*, ∀d, d^′^ ∈ D* cause side effects *r, ∀r ∈ R* is obtained by acting the sigmoid function on the score tensor 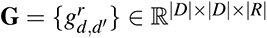 computed by the scoring function:

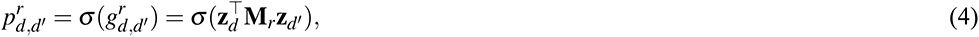

where **M***_r_* is a trainable diagonal matrix associated with the side effect *r ∈ R*. We call it a single-instance prediction or a single prediction. In addition to calculating the probability that the drug pair (*d, d^′^*) causes the side effect *r*, the proportion of variant protein related information which leads to the model to make such a prediction is also evaluated by pharmacogenomic information utilization (PIU) and protein-protein interaction information utilization (PPIU) scores:

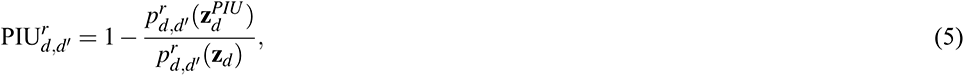

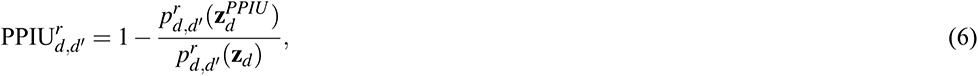

where 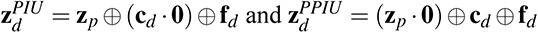.

**Algorithm 1:**
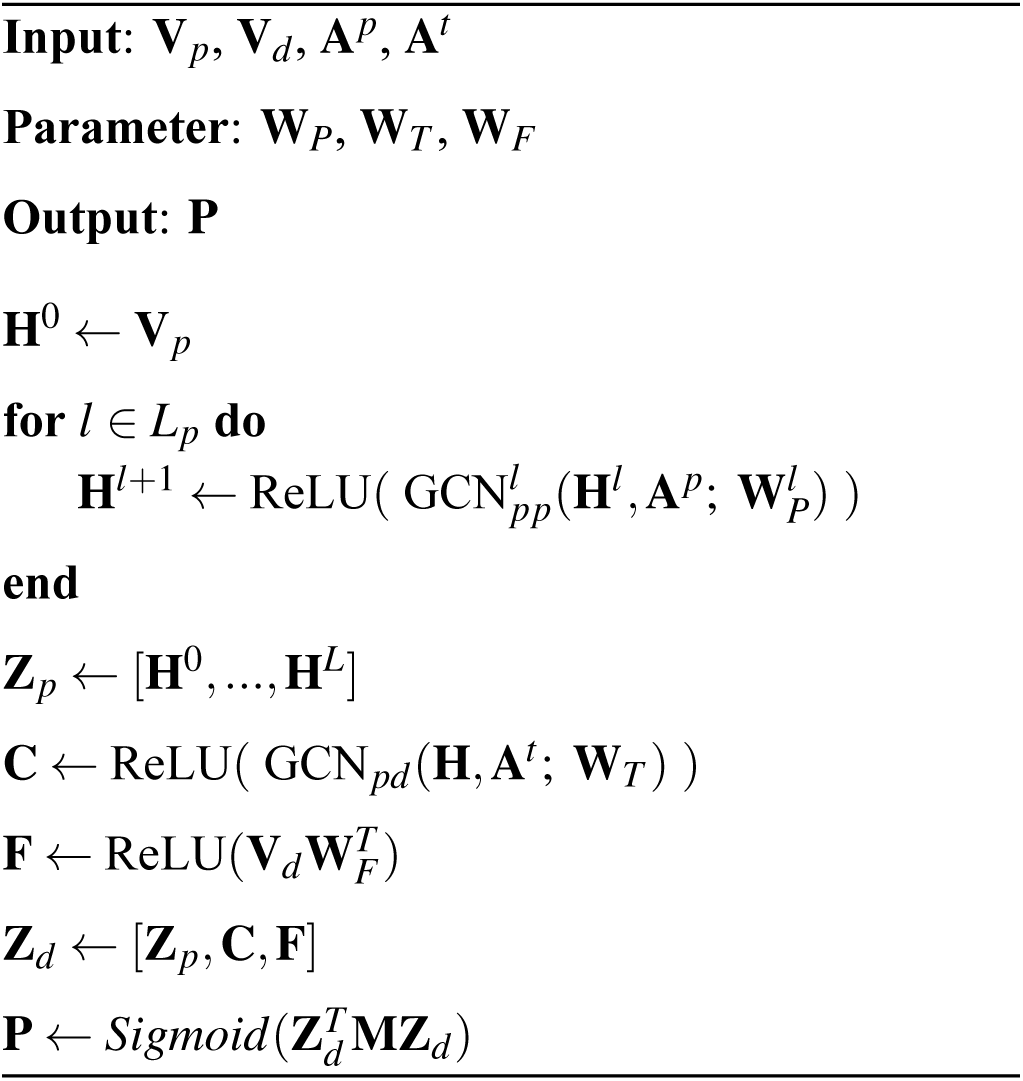
APRILE-Pred’s Forward Propagation.

**Algorithm 2:**
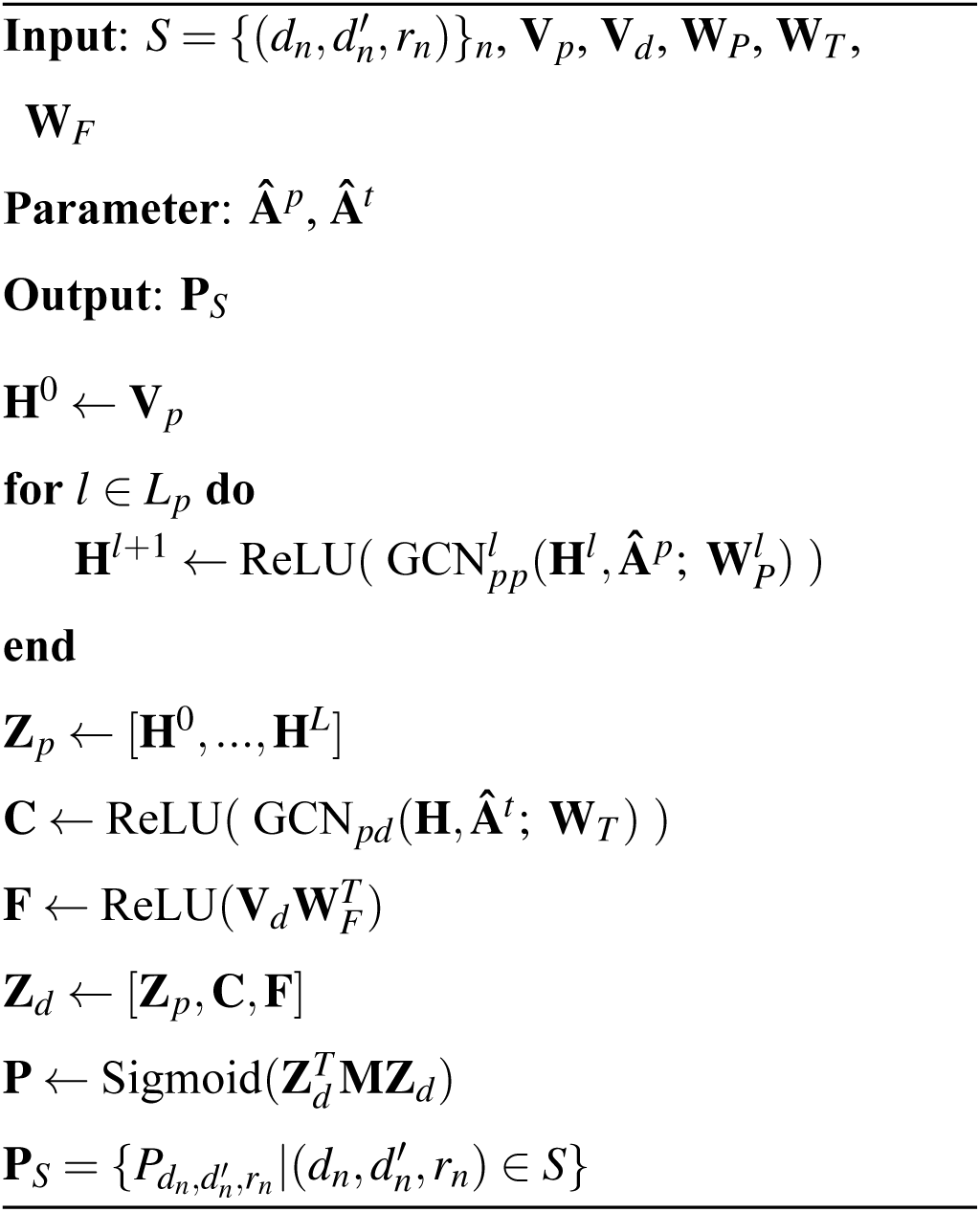
APRILE-Exp’s Forward Propagation.

### APRILE-Pred Training Details

APRILE-Pred’s forward propagation is shown in Algorithm 1. We use the Adam optimizer with a training rate of 0.01 to minimize the cross-entropy loss. We applied full-batched end-to-end training for 80 epochs on a single Tesla V100 (32 GB RAM) graphics processing unit (GPU).

#### Dataset

We use the database of polypharmacy side effects compiled by the BioSNAP-Decagon dataset^4,7^. This database consists of 964 polypharmacy side effects from 4,651,131 drug-drug interactions, which is a subset of the TWOSIDES database^45^. These polypharmacy side effects already exclude side effects attributed to an individual drug. In this work, we focus on the drugs that have at least one target protein, and the side effects that each occurred in at least 250 drug combinations. Therefore, there are 4,625,608 drug interaction edges (D-D) labeled by 861 side effects. We used as the positive polypharmacy adverse events to train and test the model.

#### Dataset split

The training and testing sets were stratified split based on side effects. To train and test the model, we need both positive and negative ADR knowledge instances in the form of (drug_1_, drug_2_, side effect). Positive instances are the observed polypharmacy side effect information, while negative ones are sampled with categorized negative sampling strategy^46^.

#### Evaluation metrics

When comparing overall performance among trained models, we use the {Micro, Macro}-averaged receiver operator characteristic area under the curve (AUROC) and the area under the precision-recall curve (AUPRC) score for all side effects. The PIU and PPIU score of each side effect are computed based on the Macro-AUROC with the same 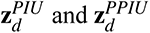 for those of each prediction.

#### Lazy training

GNNs recursively incorporate information through the information propagation paths in the graph, leading to the node intermediary embeddings capturing both graph structure and node features. However, these rich embeddings learned by APRILE-Pred prevent APRILE-Exp from finding important information sources on drug targets and non-target proteins, as the pharmacogenomic information and ADRs have been learned into drug embeddings already during the back-propagation. This makes APRILE-Pred’s ADR predictions difficult to be explained by APRILE-Exp. Therefore, to build a more interpretable model and reduce explanation redundancy for the side effects that have molecular origins, information on drug targets, protein relations and ADRs should be prevented from being learned in the trainable embedding matrices. Accordingly, we weaken the contribution of protein and drug attributes to APRILE prediction, and control the trainability of the model parameters. This strategy is referred to as ‘lazy training’. Specifically, after initializing all the model parameters by Xavier initialization^47^, we fix the parameters matrices **W**^1^_P_ (in Equation 1) and **W***_F_* (in Equation 3), which are supposed to be trainable parameters, to their initial values. The fixed parameters are referred to as lazy learners.

### APRILE-Exp: optimal explanations generation with GNNs explainer

We develop APRILE-Exp to explore the molecular mechanisms of side effects. The APRILE-Exp explains the existence of known knowledge by understanding APRILE-Pred model prediction. It is primarily motivated as an adaption of previous work on GNNExplainer^14^ for large-scale and tri-graph-like data. Given a trained APRILE-Pred model and a set of predictions we want to explain 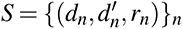, the APRILE-Exp model generates a subgraph of the original graph (the dataset) that are most influential for the APRILE-Pred model’s prediction. Therefore, the APRILE-Exp model transforms the problem of explaining why drug pairs cause side effects to finding out which protein interactions and drug interactions are the most important to predict side effects, which in turn helps us to understand the molecular mechanism of side effects.

We assigned an importance score to each edge in the P-P and P-D subgraphs by replacing the adjacency matrices **A***^p^* and **A***^t^* to **Â** *^p^* and **Â** *^t^*, where **Â** *^p^* and **Â** *^t^* have the same non-zero elements as **A***^p^* and **A***^t^* but with tunable values 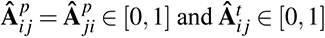. As shown in Algorithm 2, the APRILE-Exp model calculates scores of drug pairs for each side effect with the same pipeline as the prediction model but with trainable **XÂ** *^p^* and **Â** *^t^*, fixing the trained parameters **W***_P_*, **W***_T_*, and **W***_F_*. We minimize the following loss during training:

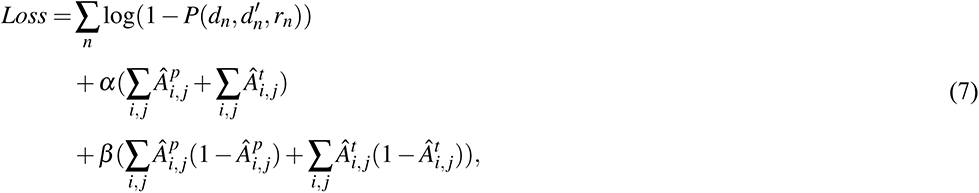

where α and β are hyper-parameters. For the loss function, the first term encourages *P* to take a value close to 1, which can be replaced with any functions that decrease sharply near 1; the second term encourages the size of subgraph given by *Â_i, j_* to be as small as possible, and it can be replaced by any monotonically increase function; and the last term = *Â_i, j_* to be either 0 or 1, which can be replaced with any function that is 0 at points *x* = 0 and *x* = 1.

### Gene Ontology Enrichment

To investigate the functional roles of the genes that are most influential for the interested predictions, we perform Gene Ontology (GO) enrichment analysis by using the GOATOOLS package^48^. The Fisher’s exact test is used to compute the p-values for each of: biological process, molecular function and cellular component, which evaluates significant enrichment of certain GO terms. We choose the non-negative Benjamini-Hochberg multiple test correction method to correct the test, which controls the false discovery rate (FDR), and use a FDR set the significance cut-off to 0.05^49^. We applied this methodology also to analyze enrichment of MeSH categories in clusters of side effects.

### Experiments

#### Lazy training sensitive study

We assessed how sensitive the model is to the initialization of these fixed matrices and the size of samples for training in detail. To this end, we trained the APRILE-Pred models with five different parameter initialization settings and six different dataset splitting rates (the proportion of positive training samples to the whole positive samples for each side effect): 0.1, 0.2, 0.4, 0.6, 0.8 and 0.9.

#### Lazy training ablation study

We trained three APRILE-Pred model 90% positive samples: with lazy training, with lazy training only on **W***_F_* (in Equation 3), and without lazy training. Model performance is tested with the remaining 10% samples and the same numbers of random negative samples.

#### Evaluation on explaining single predictions

We perform a literature-based evaluation of explanation for both learned ADR knowledge and novel predictions based on the selected trained APRILE-Pred model. To this aim, we ask APRILE-Pred to make a prediction for every drug pair and every side effect in the knowledge graph. We then divided these predictions into two groups – learned knowledge and new prediction – according to whether predictions are in the training set or not, and constructed the corresponding ranked lists based on predicted probability for each prediction group. Then, we use APRILE-Exp to obtain a subset of drug targets, protein interactions and significant GO items for each of the twenty highest ranked predictions in both rank lists, and afterwards search biomedical literature to see if we can find supporting evidence.

#### Permutation test

To assess the significance of APRILE-Exp explanations, we compared them against 10,000 random explanations (i.e., gene sets). For APRILE-Exp and random explanations, we computed the precision and Jaccard index using the CTD^17^ disease-associated genes as validation. We compared two types of explanations: all genes in a APRILE-Exp subgraph, or the subset of these genes that are in enriched GO terms. Additionally, two types of random models were tested: (A) one that preserves the number of drug target vs. non-target genes, and (B) one that samples from all genes in the KG. Permutation test p-values were determined as the number of random explanations whose precision (or Jaccard index) was equal to or greater than that of the APRILE-Exp explanation.

## Acknowledgements

This work was supported by Queen’s University, and the Natural Sciences and Engineering Research Council of Canada (NSERC) [RGPIN-2020-06325]. This research was enabled in part by support provided by Compute Ontario (www.computeontario.ca) and Compute Canada (www.computecanada.ca). Computations were performed on resources and with support provided by the Centre for Advanced Computing (CAC) at Queen’s University in Kingston, Ontario. The CAC is funded by: the Canada Foundation for Innovation, the Government of Ontario, and Queen’s University.

## Author contributions

LY and HX conceived the study. HX and SS developed the framework. HX, LY, HY and AIH carried out the simulations and biologically interpreted results. All authors wrote and edited the manuscript.

## Data availability

The data for constructing pharmacogenomic knowledge graph and drug-drug interactions is available in the BioSNAP^7^ database (http://snap.stanford.edu/biodata/). The medical subject headings for drugs are available at https://meshb.nlm.nih.gov. The CTD^17^ disease-gene associations are available in http://ctdbase.org/downloads/.

## Code availability

Code to train, test and select an APRILE-Pred model is available at https://github.com/QCSB/APRILE-Pred. Code to build and use APRILE-Exp is available at https://github.com/QCSB/APRILE-Exp. The pub-lished APRILE python package is available at https://github.com/QCSB/APRILE

## Supplementary information

The supplementary information includes four figures and three tables. The supplementary data is available at https://github.com/QCSB/APRILE-Exp/tree/master/supplementary-data.

## Supplementary Information

### Supplementary Note

#### Predicted explanations offer novel mechanistic hypotheses

A patient’s existing conditions may influence the mechanism of adverse reactions. We investigated one such example in which the combination of glipizide and lansoprazole may result in hypophosphatemia. APRILE predicted the likelihood of this side effect at 97% .

Glipizide is a second-generation oral hypoglycemic agent used to treat type 2 diabetes that targets the *CYP* gene family^1^. Lansoprazole ia a proton pump inhibitor commonly used to reduce gastric acid and treat gastric ulcers^2^ and is metabolized by the liver using the P450 pathway; its targets include *PHOS-PHO1*, a gene involved in the production of inorganic phosphate from organic compounds. Lansoprazole inhibits *PHOSPHO1*, disturbing this phosphorous cycle, which causes a drop in serum phosphate level. Meanwhile, Glipizide inhibits *CYP3A4*—a member of the P450 pathway—causing a delayed excretion and increased exposure to lansoprazole^3^, greatly reducing the amount of serum phosphate and ultimately leading to hypophosphatemia. Glipizide also inhibits *ABCC8* from promoting insulin production by the liver^1^, thus having the same effect as direct insulin therapies. Insulin therapies may lead to a redistribution of phosphate due to insulin-stimulated cells actively acquiring phosphate from the blood^4^. This mechanism is not sufficient to cause hypophosphatemia but may worsen an existing condition^5^. Glipizide is used to treat type 2 diabetes, and these patients have more renal phosphate excretion than average due to osmotic diuresis^6^. This condition is a potential contributing factor to an existing hypophosphatemia adverse effect. Thus, APRILE was able to offer a comprehensive explanation for hypophosphatemia arising from the combination of glipizide and lansoprazole (Figure S4).

---

**Figure S1.**
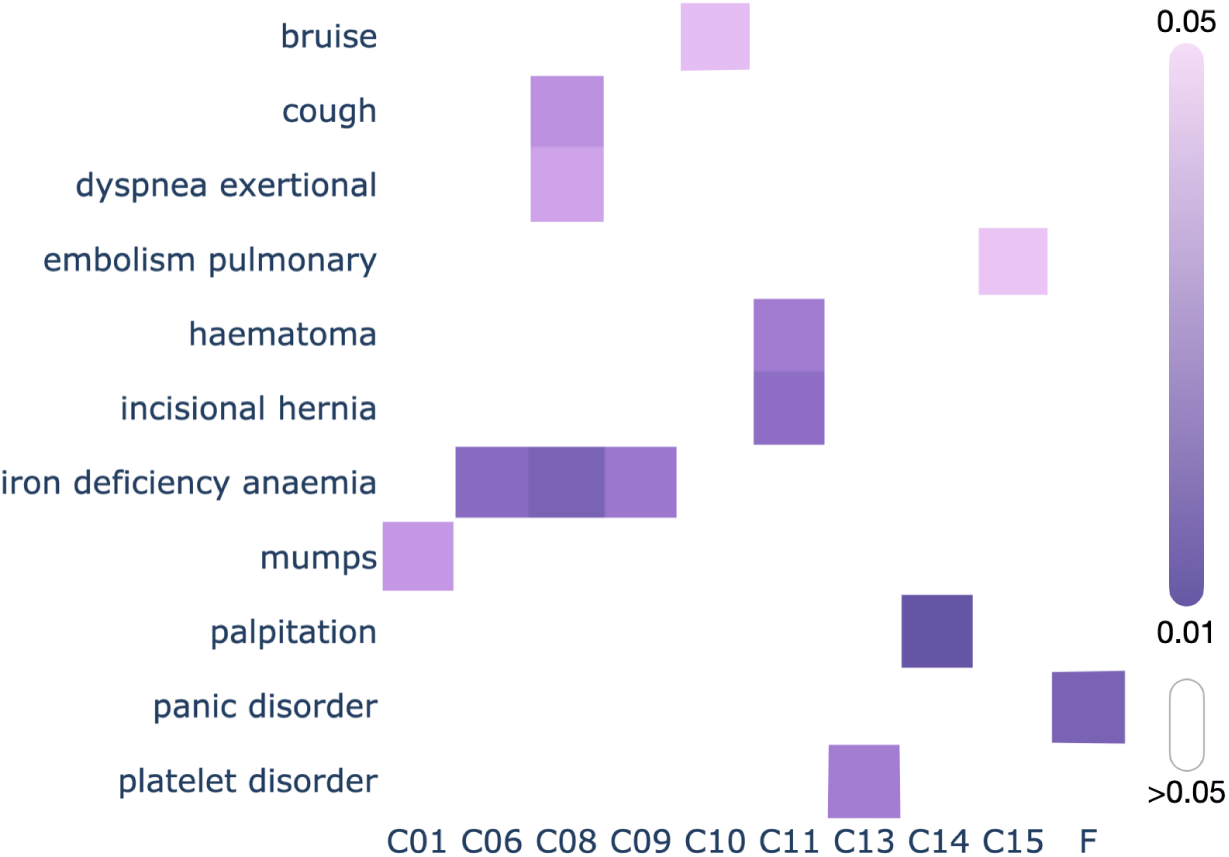
Significant relations (*p <* 0.05) among side effect clusters and MeSH disease categories.

**Figure S2.**
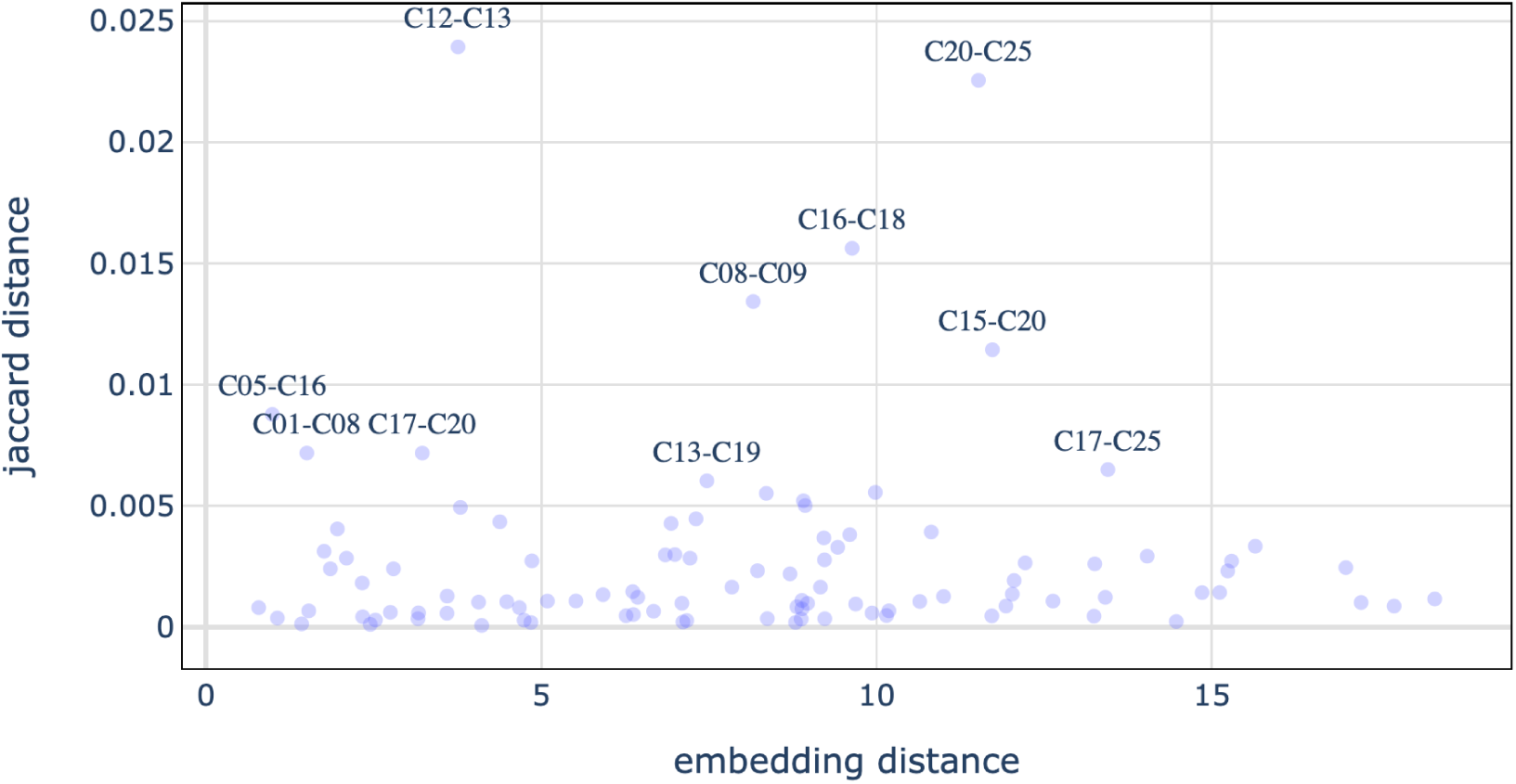
Jaccard distance between side effects vs. Euclidean distance between embeddings learned by APRILE-Pred for each pair of categories (22 in total) in Fig. 2d. The markers are shown if their Jaccard distance *>* 0. The labels of markers are shown if their Jaccard distance *>* 0.006.

**Figure S3.**
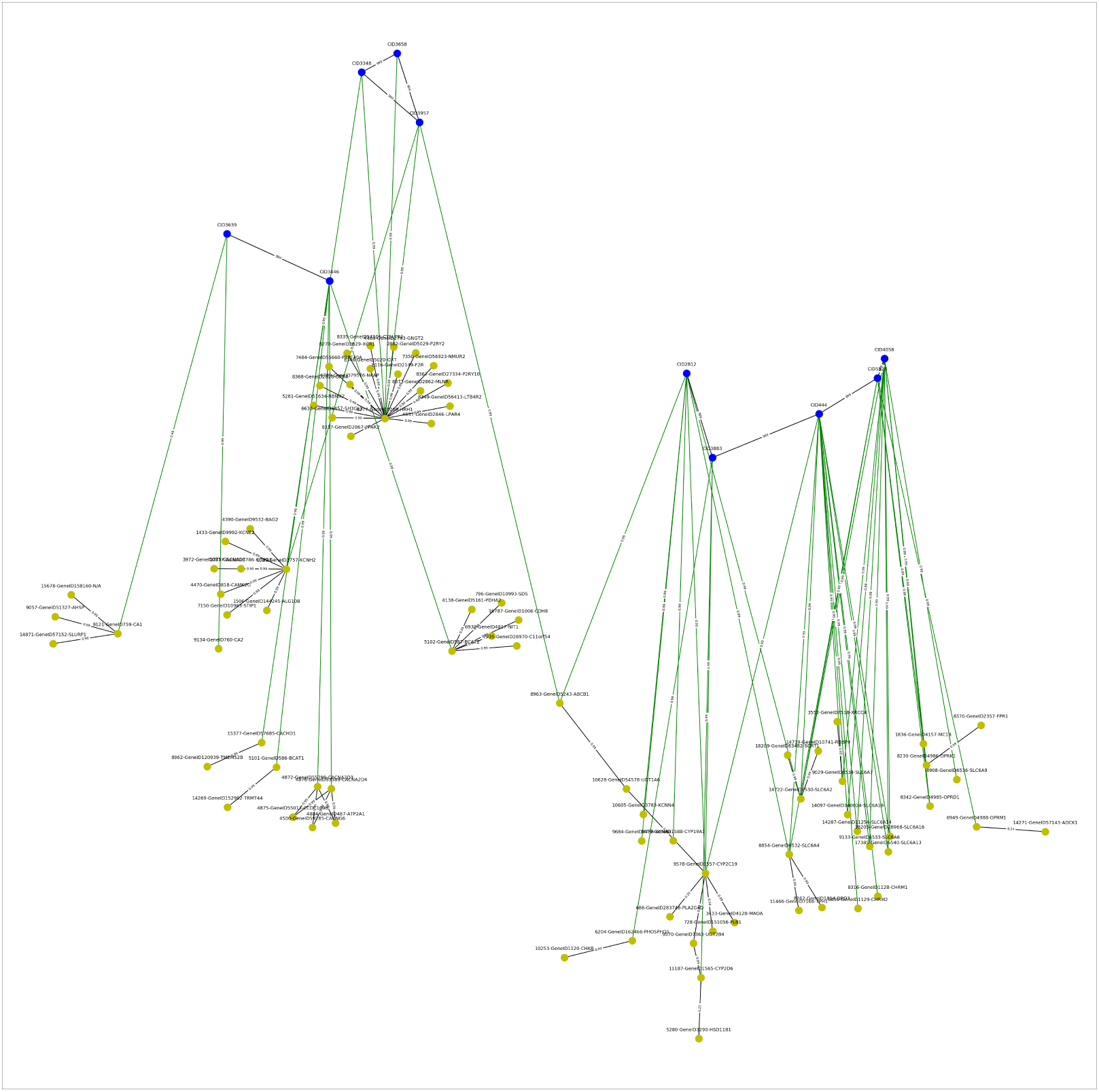
APRILE explanation for panic disorder caused by drug pairs (10 possible drugs). Parameters used: regulation weight of 2.0, and probability threshold of 0.99. Drug nodes are blue and protein nodes are yellow.

**Figure S4.**
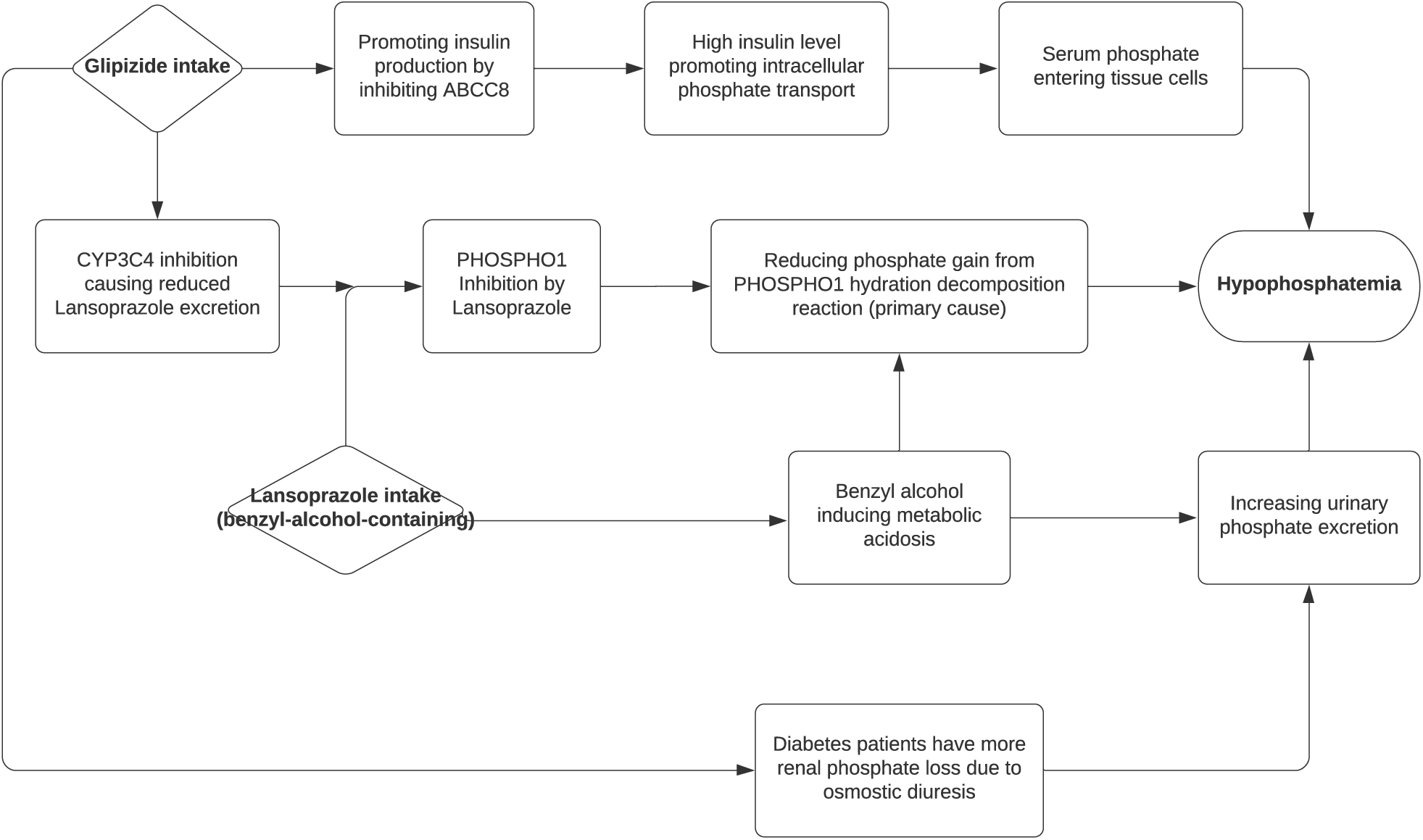
Hypothesized mechanism for hypophosphatemia caused by glipizide-lansoprazole interaction.

**Table S1.**
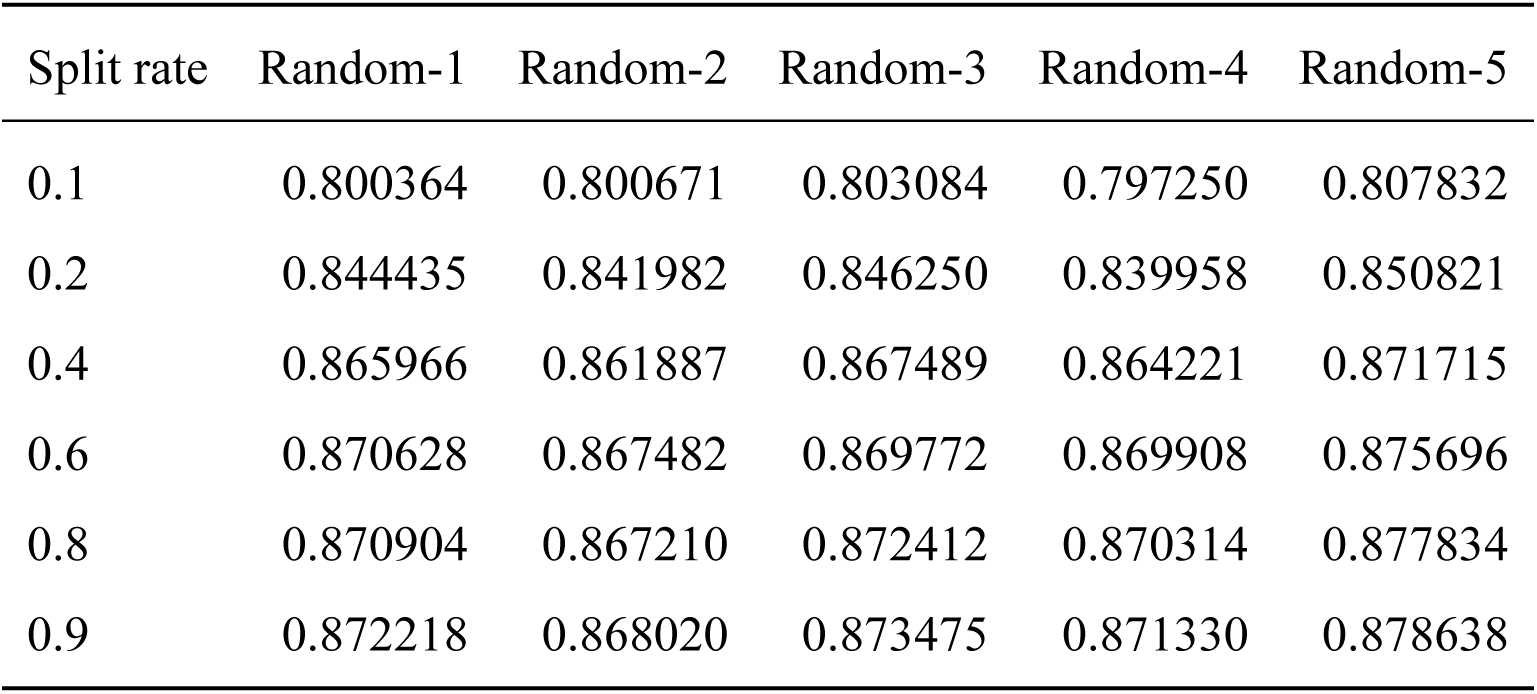
Macro-averaged AUROC for applying lazy training with different split rates and random states for parameter initialization

| Split rate | Random-1 | Random-2 | Random-3 | Random-4 | Random-5 |
| --- | --- | --- | --- | --- | --- |
| 0.1 | 0.800364 | 0.800671 | 0.803084 | 0.797250 | 0.807832 |
| 0.2 | 0.844435 | 0.841982 | 0.846250 | 0.839958 | 0.850821 |
| 0.4 | 0.865966 | 0.861887 | 0.867489 | 0.864221 | 0.871715 |
| 0.6 | 0.870628 | 0.867482 | 0.869772 | 0.869908 | 0.875696 |
| 0.8 | 0.870904 | 0.867210 | 0.872412 | 0.870314 | 0.877834 |
| 0.9 | 0.872218 | 0.868020 | 0.873475 | 0.871330 | 0.878638 |

**Table S2.** Literature evidence for APRILE-Exp explanations for training data.

| Side effect | Drug target? | Gene | Supported | Literature evidence |
| --- | --- | --- | --- | --- |
| scotoma | target | PDE6B | yes | PDE6B mutations cause Retinitis Pigmentosa (RP)<sup>7</sup>. Scotoma is a symptom for RP patients. The polypharmacy may affect PDE6B similar to mutations. |
| scotoma | target | PDE6C | yes | PDE6C mutations cause achromatopsia<sup>8</sup>. Scotoma is a symptom of achromatopsia. |
| elevated prostate specific antigen | target | PDE11A | no | — |
| melanocytic naevus | non-target | CD7 | yes | Patient with melanocytic nevus exhibited low CD7 expression levels<sup>9,10</sup> |
| bacterial pneumonia | target | CA4 | indirect | CA4 is expressed in the lung and kidney and its inhibition alters CO<sub>2</sub> equilibrium in tissues<sup>11</sup>. In lambs with pneumonia, an insignificant decrease of carbonic anhydrase activity was observed<sup>12</sup>. |
| post traumatic stress disorder (PTSD) | target | PDE6B | indirect | No direct evidence but PDE6B was differentially expressed in hippocampi of alcoholics, potentially affecting cortisol pathway dysregulation, which is observed in PTSD<sup>13</sup>. |

**Table S3.**
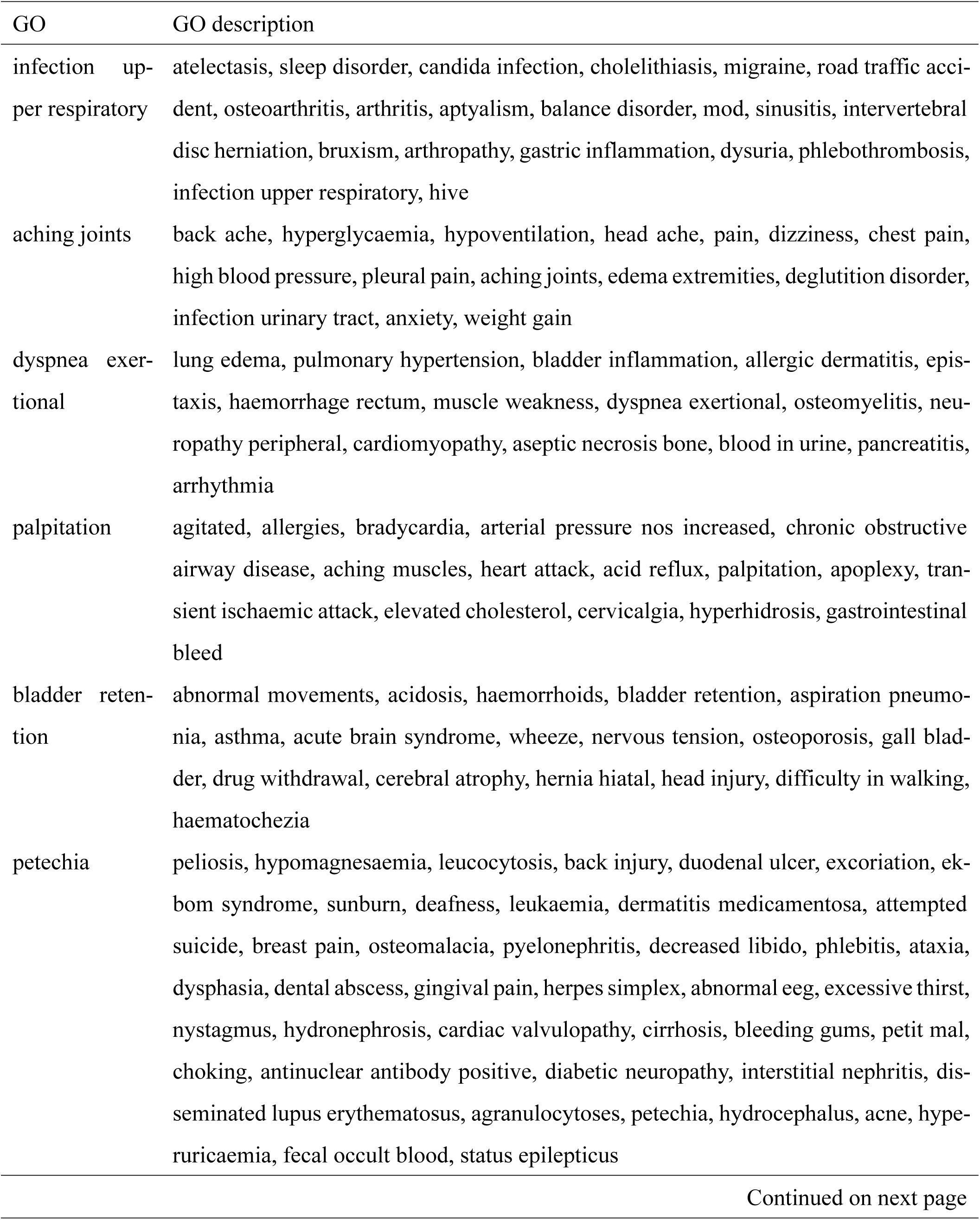
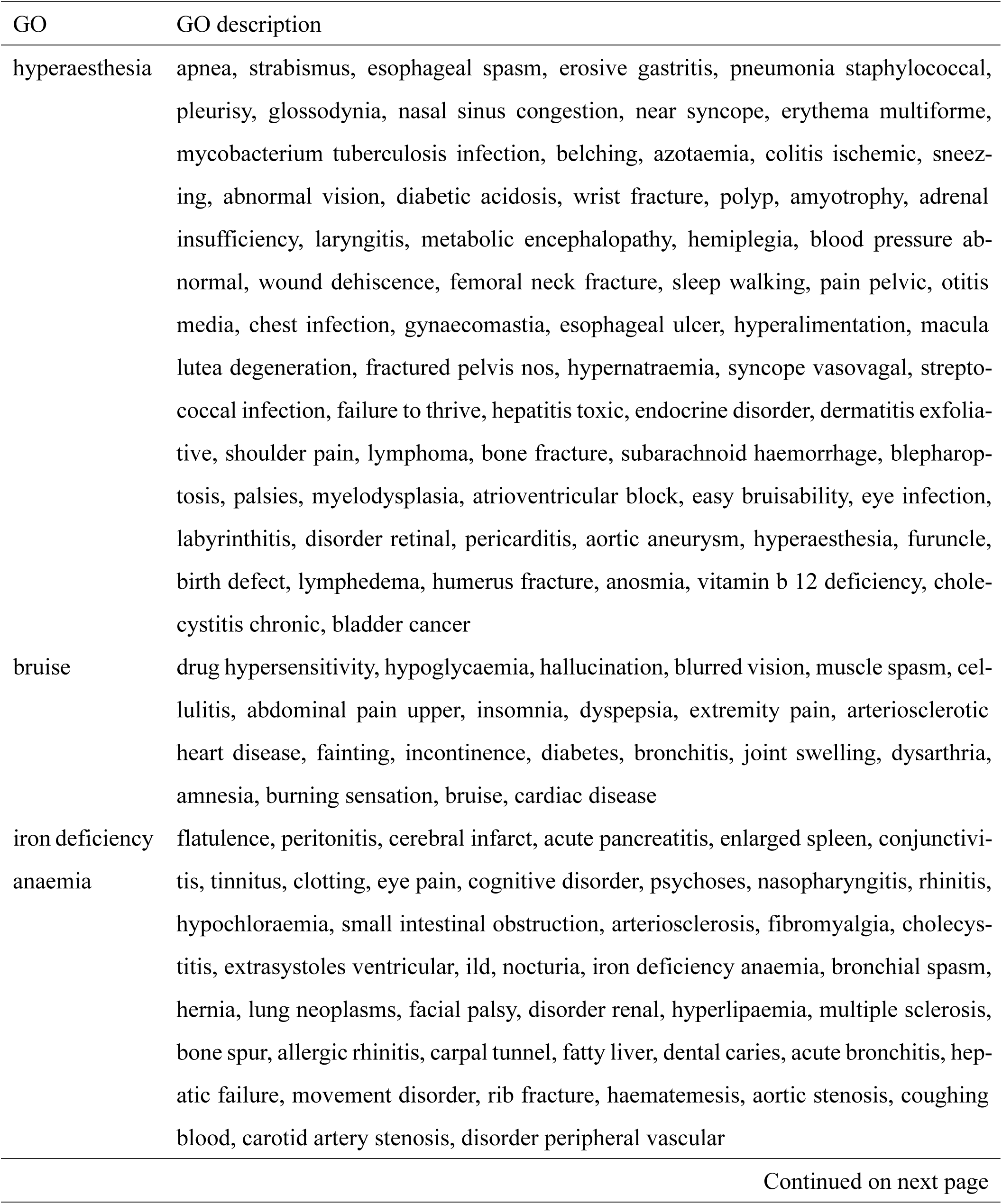
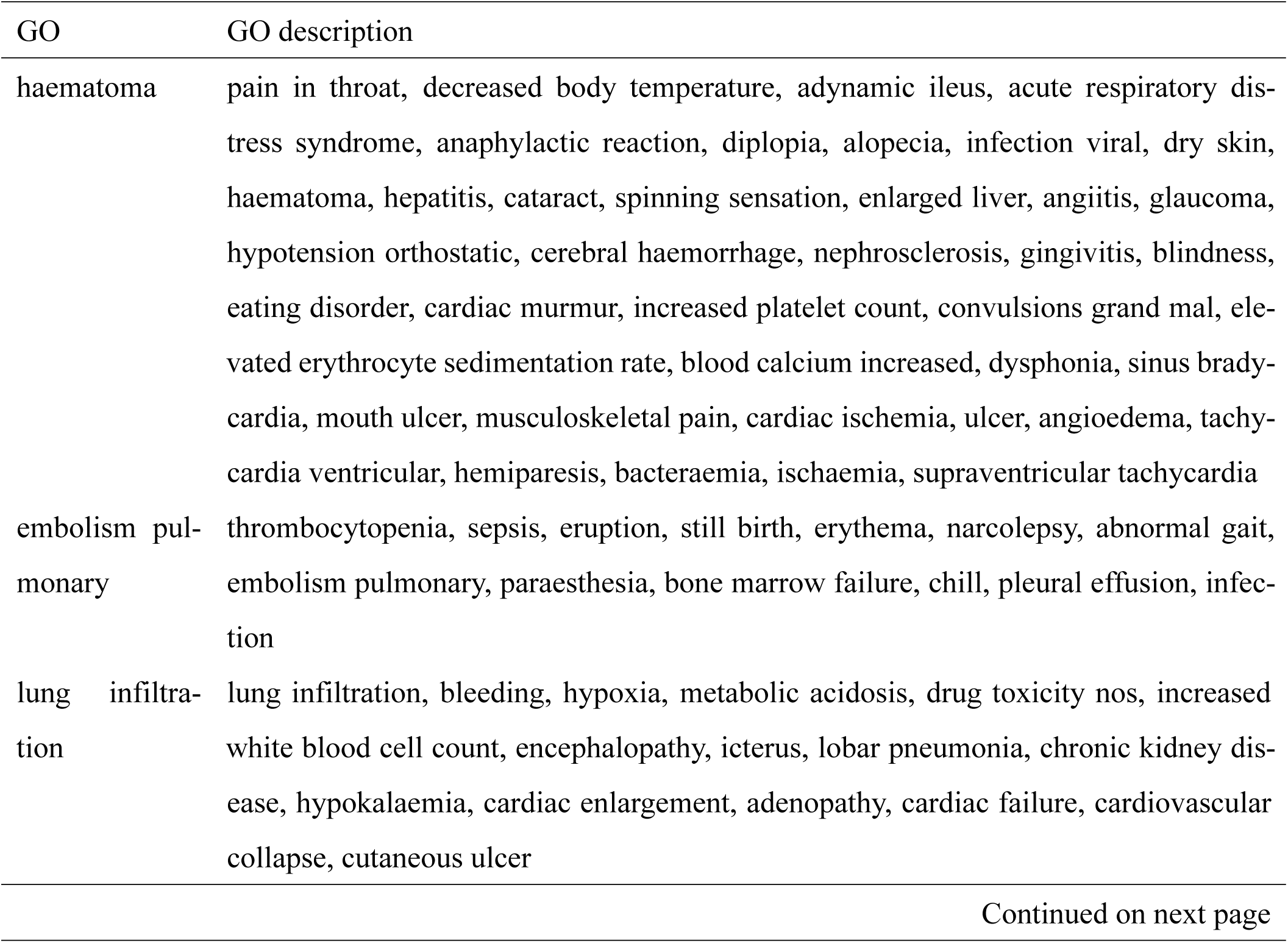
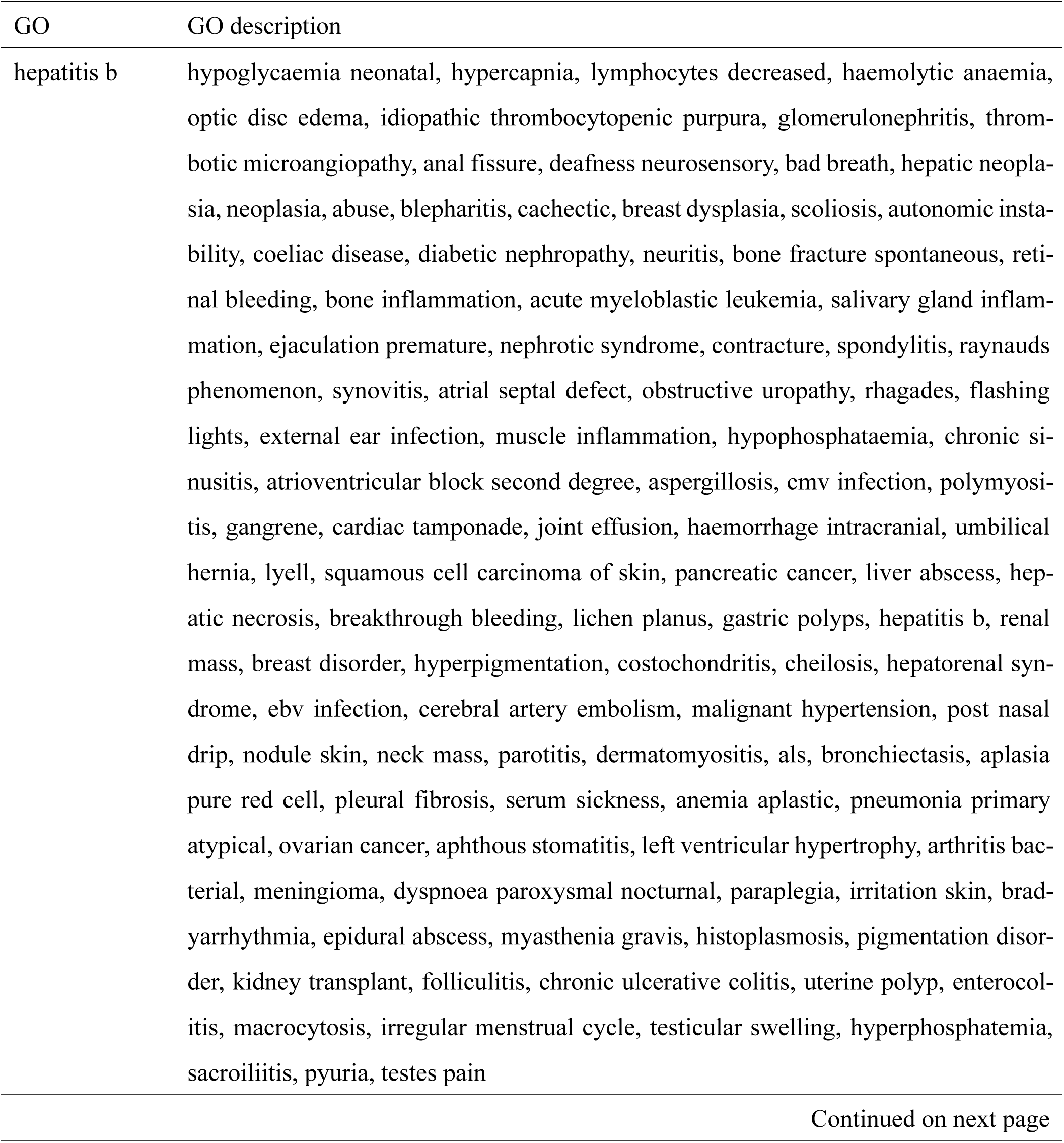
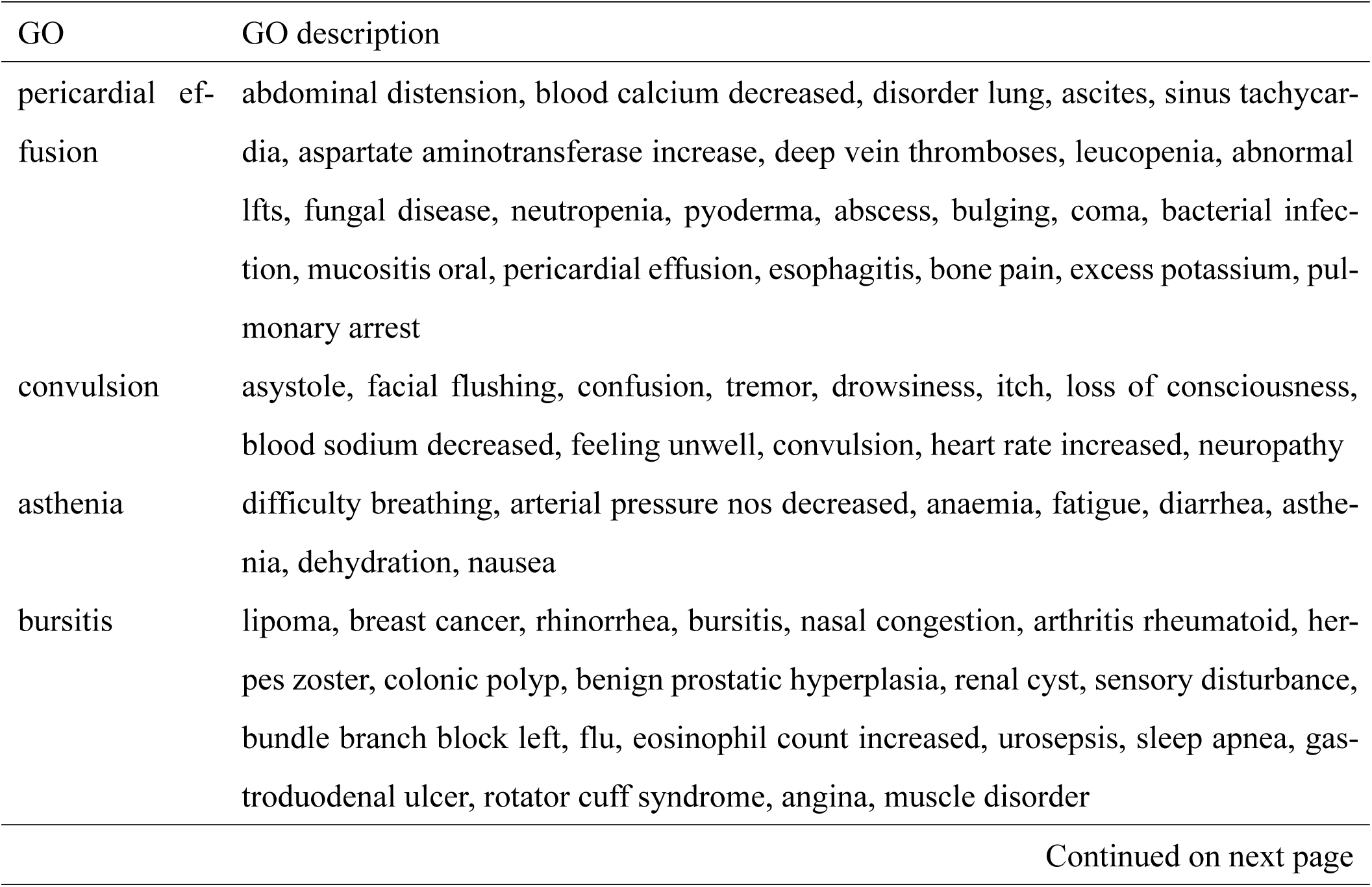
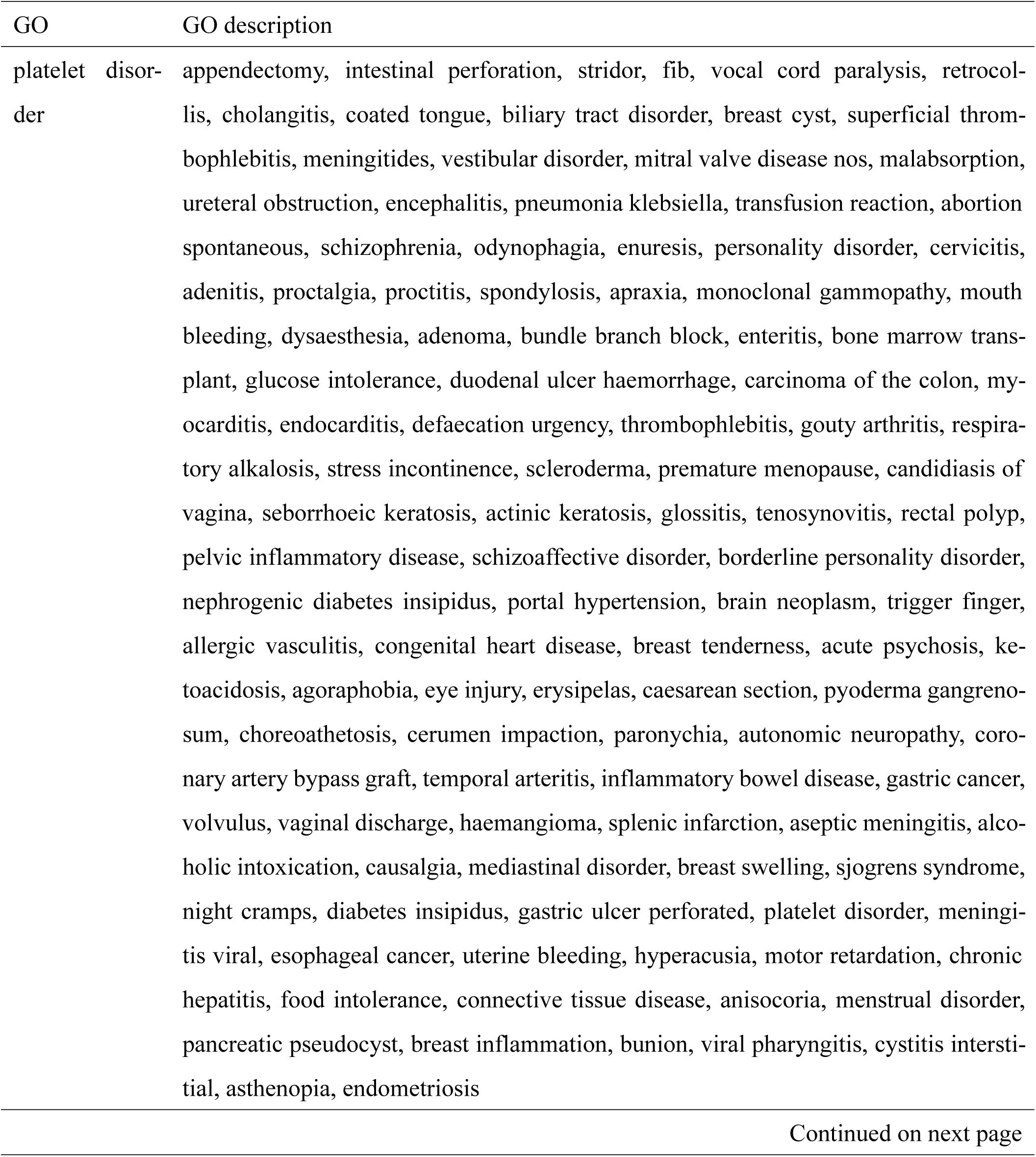
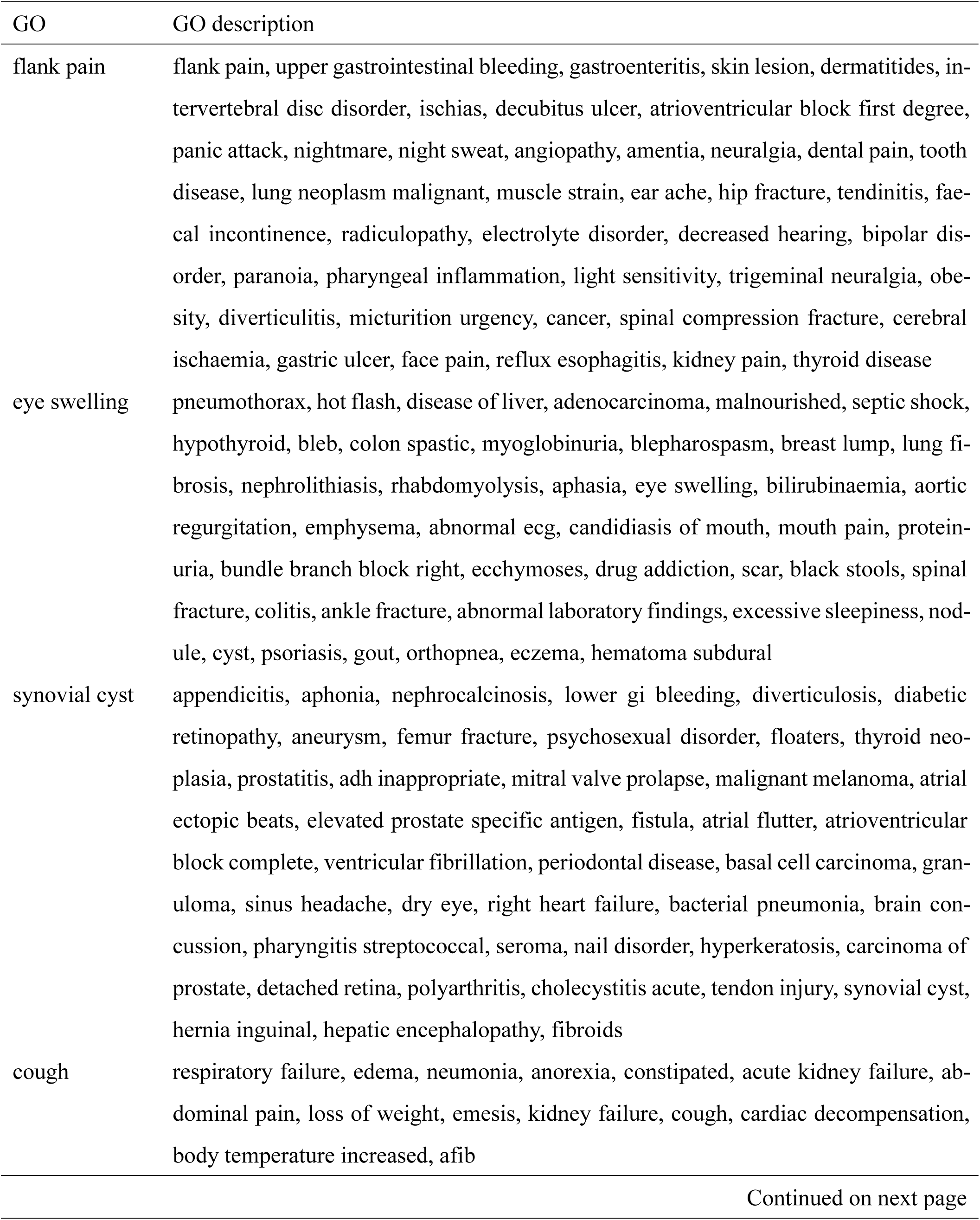
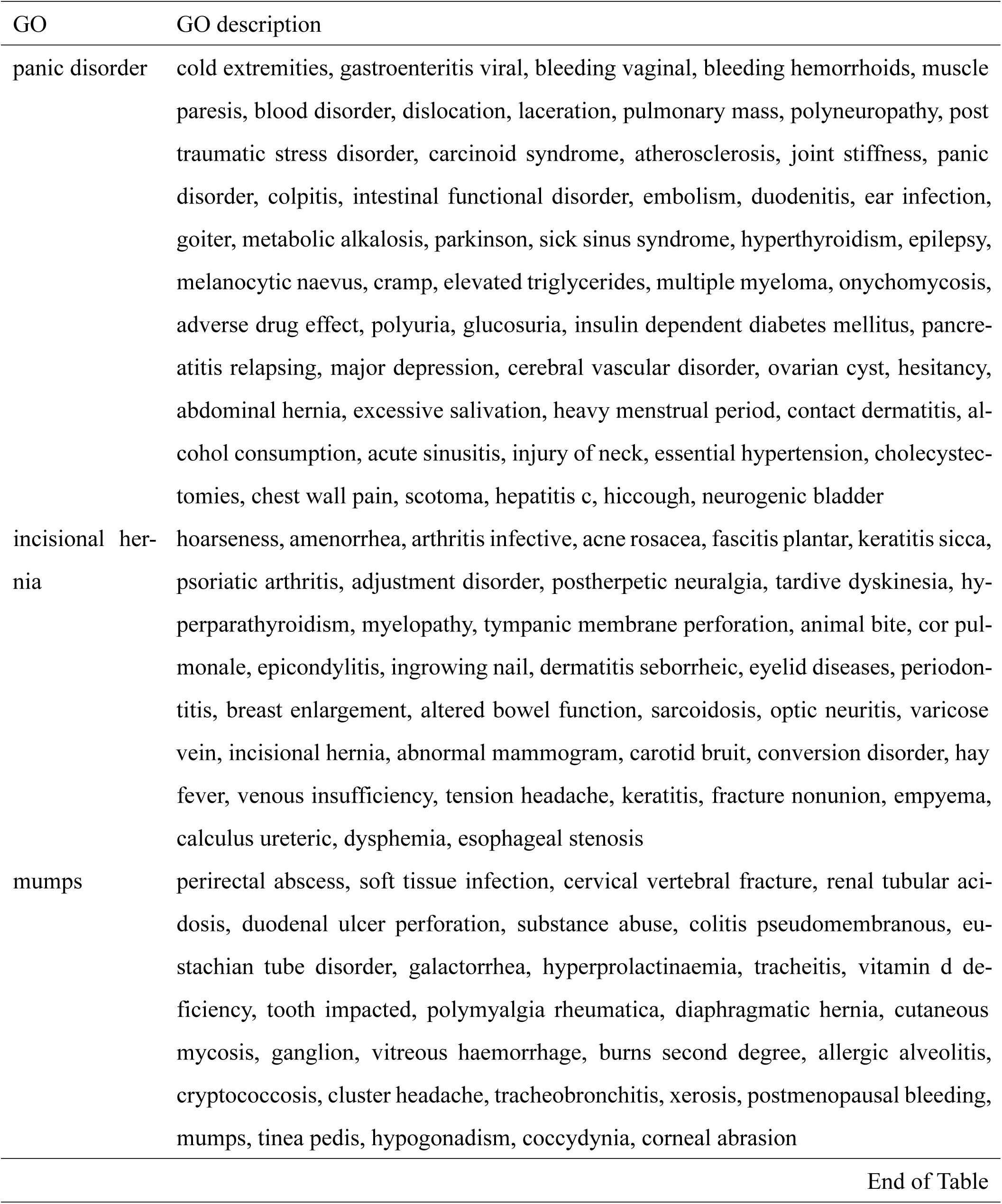
Side effect clustering based on the identified biological processes by APRILE.

**Table S4.** Commonly enriched GO terms identified by APRILE-Exp for three peptic ulcers

| GO | GO description | Literature evidence |
| --- | --- | --- |
| GO:0045540 | Regulation of cholesterol biosynthetic process. | Cholesterol increases the resistance of *H. pylori*, and the clinical manifestations of this bacteria include peptic ulcer disease<sup>14</sup>. *Helicobacter pylori* is associated with dyslipidemia but not with other risk factors of cardiovascular disease. |
| GO:0055114 | Oxidation-reduction process. | Derangements of redox homeostasis in the gastrointestinal tract are believed to lead to peptic ulcer<sup>15</sup>. |
| GO:0019369 | Arachidonic acid metabolic process. | Arachidonic acid metabolites play an important role in gastrointestinal homeostasis and have a significant impact in preventing or treating peptic ulcers<sup>16</sup>. |
| GO:0033384 | Geranyl diphosphate biosynthetic process. | Farnesyl diphosphate and geranyl diphosphate are both affected by the direct interaction between bisphosphonates and a biosynthetic enzyme<sup>17</sup>. Bisphosphonates are drugs used as treatment for osteoporosis that can cause serious inflammation and ulcers in the GI tract<sup>18</sup>. |
| GO:0006805 | Xenobiotic metabolic process. | Xenobiotics are widely used in the treatment of Parkinson disease, and Parkinson patients under treatment are prone to develop peptic ulcers, hence it is thought that there is a correlation<sup>19</sup>. |
| GO:0005789 | Endoplasmic reticulum membrane. | Research suggests that the endoplasmic reticulum would be the main source for autophagosomes' membrane, which is formed for autophagic processes during an ulcerative infection<sup>20</sup>. |

Table S4 – continued from previous page
| GO | GO description | Literature evidence |
| --- | --- | --- |
| GO:0020037 | Heme binding. | Some proteins that are involved in heme binding are thought to be one important virulence factor of *H. pylori*<sup>21</sup>. |
| GO:0005506 | Iron ion binding. | Iron ion binding capacity is thought to be lower in people with *H. pylori* infection<sup>22</sup>. |
| GO:0102568 | Phospholipase A2 activity consuming 1,2-dioleoylphosphatidylethanolamine. | This is directly correlated to the arachidonic acid metabolic process and to gastrointestinal ulceration<sup>23</sup>. |
| GO:0004623 | Phospholipase A2 activity. | Both this term and the previous one refer to the activity of phospholipase A2, which is a mucosal barrier breaker in peptic ulcer disease<sup>24</sup>. |
| GO:0008395 | Steroid hydroxylase activity. | Steroid hydroxylase is involved in steroid biosynthesis and a too high concentration of steroid can lead to gastrointestinal ulcers and bleeding<sup>25</sup>. |
| End of Table |  |  |

